# Phenological segregation suggests speciation by time in the planktonic diatom *Pseudo-nitzschia allochrona* sp. nov.

**DOI:** 10.1101/2021.05.28.446110

**Authors:** Isabella Percopo, Maria Valeria Ruggiero, Diana Sarno, Lorenzo Longobardi, Rachele Rossi, Roberta Piredda, Adriana Zingone

## Abstract

The emergence of new species is poorly understood in marine plankton, where the lack of physical barriers and homogeneous environmental conditions limit spatial and ecological segregation. Here we combine molecular and ecological information from a long term time series and propose *Pseudo-nitzschia allochrona*, a new cryptic diatom, as a possible case of speciation by temporal segregation. The new species differs in several genetic markers (18S, LSU and ITS rDNA fragments and *rbc*L) and is reproductively isolated from its closest relative, which is morphologically identical. Data from a long term plankton time series show *Pseudo-nitzschia allochrona* invariably occurring in summer-autumn in the Gulf of Naples, where its sister species are instead found in winter-spring. Temperature and nutrients are the main factors associated with the occurrence of *P. allochrona*, which could have evolved in sympatry by switching its phenology and occupying a new ecological niche. This case of possible speciation by time shows the relevance of combining ecological time series with molecular information to shed light on the eco-evolutionary dynamics of marine microorganisms.

## Introduction

The mechanisms underlying the emergence of new microbial species in the marine realm are poorly explored. In the plankton, the diversity of such a relevant group of organisms has long been underestimated because of the scarcity of morphological features and the lack of adequate tools for the discrimination of meaningful units of ecology and evolution. The advent of molecular approaches in taxonomy and ecology has revolutionized our perception of unicellular microalgae diversity, revealing consistent genetic diversity coupled with morphological stasis in many instances and leading to an escalation in the discovery of cryptic or pseudocryptic species. The evidence of a so far hidden diversity in marine microbes has increasingly emerged with the discovery of multiple species within iconic taxa long considered to be a single one. Notable examples are the diatoms *Skeletonema* ^1,2^ and *Leptocylindrus* ^3^ and the prasinophyte *Micromonas* ^4^. Even higher levels of diversity have emerged from massive sequencing of environmental DNA, which has revealed high level of interspecific and intraspecific genetic diversity in the microbial realm ^5,6^.

Marked eco-physiological differences and distinct biogeographic patterns among pseudocryptic and cryptic species definitely indicate the functional value of their diversity in plankton ecology. At the same time, the discovery of the coexistence of many hardly distinguishable organisms in an apparently homogeneous environment exacerbates the so called ‘paradox of plankton’ ^7^, based on the idea that competitive exclusion in such a resource-limited environment as the ocean should favor few fittest species occupying large, unstructured niches. At the global scale, genetic differences within taxa previously considered ubiquitous challenge the Baas-Baker postulate of a limited number of widespread microbes – namely, the idea that ‘*everything is everywhere*’ ^8^.

The question left open by the finding of the last decades is how this great diversity may arise at sea, and particularly in the plankton, where physical barriers are virtually absent and the species’ dispersal potential is unlimited. In these conditions, the role of geographic separation in promoting species diversification in allopatry appears unlikely ^9^. Sympatric speciation driven by ecological segregation ^10,11^ could also be limited for planktonic microalgae in the often remixed photic zone. In the terrestrial habitat, the divergence of breeding times, i.e., allochronic segregation, has been posited as a plausible mechanism for sympatric speciation, whereby phenological changes in part of the population can promote assortative mating and genetic divergence between subpopulations ^12,13^. These possibility has rarely been considered for aquatic organisms ^14^ and never for unicellular algae.

The pennate diatoms *Pseudo-nitzschia* are needle-like, chain-forming planktonic organisms thriving in coastal waters around the world’s seas. Their life cycle includes heterotallic sexual reproduction, through which these species re-establish maximal cell size ^15^. The diversity of the genus, now including 58 species ^16^, has expanded over the years due to the raised attention to the production in some species of neurotoxins (domoic acid) that cause a syndrome known as amnesic shellfish poisoning. The use of molecular markers has much contributed to the description of new species within species-complexes of taxa that are hardly distinguishable morphologically, not even with electron microscopy. These cryptic and pseudocryptic species may show distinct geographic ranges and temporal patterns ^17,18^, as well as different biochemical and functional traits ^19^, while the intricacy of phylogenetic patterns could be the result of cases of introgression and speciation by recombination ^20^.

In the Gulf of Naples (GoN) twelve *Pseudo-nitzschia* species are recorded ^18,21^. Among them, the *P. delicatissima*-complex ^22^ is the most represented group, with three different species only identifiable by means of molecular methods: *P. delicatissima* sensu stricto, *P. arenysensis* and *P. dolorosa*. Here we describe another cryptic *P. delicatissima*-like species as *P. allochrona* sp. nov. based on sequences of diagnostic nuclear (18SrDNA, 28SrDNA, ITSrDNA) and chloroplast (partial RUBISCO, *rbc*L) markers, ITS2 secondary structure and interbreeding experiments with its sibling species *P. arenysensis*. By coupling molecular information with environmental data from a 30 ys-long time series, we build on the phenological and ecological peculiarities of *P. allochrona* to discuss possible mechanisms of speciation in the plankton realm.

## Materials and methods

### Samples and cultures

Sixty-two strains of *Pseudo-nitzschia allochrona* sp. nov. were isolated from surface waters of the Gulf of Naples (Mediterranean Sea) from 2007 to 2016, mainly from the Long Term Ecological Research Station MareChiara (LTER-MC, 40° 48.5’ N, 14° 15’ E, depth ca 75 m) ^23,24^ (Suppl. Table 1). Two additional strains were isolated from the Ionian Sea (Mediterranean Sea) in September 2008. In addition, in this study we considered 187 strains of other *P. delicatissima*-like species obtained from the Gulf of Naples, which had been identified as *P. arenysensis*, *P. delicatissima* or *P. dolorosa* with molecular analyses in previous studies ^25–27^, for which we could track the isolation date. Two further strains of *P. arenysensis* from the Gulf of Naples (BB16 and CM63, courtesy of M. Ferrante, SZN) were used in mating experiments. All strains were isolated by hand pipetting. Cultures were grown in F/2 medium, and maintained at 20°C under an irradiance of 70-80 μmol photon m^−2^ s^−1^ and a 12:12 light:dark regime.

### Microscopy

Live culture material of *P. allochrona* was observed and cell measurements taken under a Zeiss Axiophot and an Axiovert 200 light microscopes (Carl Zeiss, Oberkochen, Germany). Pictures were taken with a Zeiss Axiocam digital camera (Carl Zeiss, Oberkochen, Germany). For transmission electron microscopy (TEM) observations, clean diatom frustules were obtained boiling culture material for a few seconds with nitric and sulphuric acids (1:1:4, sample:HNO_3_:H2SO_4_) to remove organic matter, and washing it with distilled water (modified from ^28^). The material was then mounted on Formvar-coated grids and observed with a Philips 400 TEM (Philips Electron Optics BV, Eindhoven, Netherlands). For scanning electron microscopy (SEM), material from successful mating experiments was fixed with glutaraldehyde (final concentration 2.5% v/v), placed on a filter in a Sweenex filter holder and dehydrated in a graded ethanol series (30-100%). Filters were critical-point dried, mounted on stubs, sputter-coated with gold-palladium and observed with a JEOL JSM-6500F SEM (JEOL-USA Inc., Peabody, MA, USA).

### Toxin analysis

Cultures of *P. allochrona* strains SZN-B495 and SZN-B524 were grown in 1 l Erlenmeyer flasks (20 °C, irradiance 70-80 μmol photon m^−2^ s^−1^ and 12:12 light:dark regime), harvested at their late exponential growth phase and centrifuged (2,000 rpm for 10 min). The cell pellet was stored at −18 °C until analysis. Cells were lysed by sonication for 5 min, then added with 500 μl of MeOH/H_2_O (1:1) mix and vortexed for 3 min. Cells were lysed by sonication again for 5 min and centrifuged at 3,000 rpm for 5 min. The clear supernatant was transferred into a glass test tube and the pellets were resuspended in 500 μl of MeOH/H_2_O (1:1) and vortexed for 3 min. All steps were repeated 3 times. Then the supernatant was evaporated and the residue resuspended with 500 μl of MeOH/H_2_O (1:1). This solution was centrifuged at 9,000 rpm for 5 min and finally 5 μl were analyzed by LC-MS/TOF. Certified standard of domoic acid was purchased from the National Research Council of Canada (NRCC). Acetonitrile, methanol and water were HPLC grade. Trifluoroacetic acid was obtained from VWR International (USA).The LC consisted of an Agilent 1100 instruments equipped with binary pump and an auto sampler. Phenomenex Luna 3μ PFP(2) (150 x 2.00 mm) was used for chromatographic separation. The isocratic mobile phase consisted of a mixture of 0.02% aqueous trifluoroacetic acid and acetonitrile in the ratio 90:10 (v/v), isocratic elution of 10% B at 0-15 min. The flow rate was 0.2 ml/min. Sample solutions (8, 4, 2, 0.4 ppm) were prepared in ACN/W (1:9) and 5 μl was injected. The MS/TOF analysis worked in positive ion mode, and mass range was set at *m/z* 100-1,000 u at a resolving power of 10,000. The conditions of ESI source were as follows: drying gas (N_2_) flow rate, 11 ml/min; drying gas temperature, 350°C; nebulizer, 45 psig; capillary voltage, 4,000 V; fragmentor 225 V; skimmer voltage, 60 V. All the acquisition and analysis of data were controlled by Agilent LC-MS TOF Software (Agilent, USA-Germany). Tuning mix (G1969-85001) was used for lock mass calibration in our assay. Under these conditions, major peaks of DA would appear as the protonated ion at *m/z* 312, being accompanied by minor peaks consisting of sodium-binding ions at *m/z* 334. For the reference DA material, LOD is 0.001μg/kg (1 ng) and LOQ 0.01μg/kg (10 ng).

### Molecular analyses

Exponentially growing cultures were filtered on 0.8 μm pore size Isopore membrane filters (Millipore, Schwalbach, Germany). Genomic DNA was extracted using the CTAB buffer as in ^29^ and used as a template for the amplification of the following loci: partial 18S rDNA using Euk-A and Euk-B primers ^30^; hypervariable (D1/D2) 28S rDNA region using DIR and D3Ca primers ^31^; ITS-rDNA using ITS-1 and ITS-4 primers ^32^; partial RUBISCO (*rb*cL) using *rbc*L1 and *rbc*L7 primers ^25^. PCRs were carried out in a PTC-200 Peltier Thermal Cycler (MJ Research, San Francisco, CA, USA) using reaction conditions as in the above cited references for each of the amplified loci. The amplified fragments were purified using a QIAquick PCR purification kit (Qiagen Genomics, Bothell, WA, USA) following the manufacturer‘s instructions, sequenced with the BigDye Terminator Cycle Sequencing technology (Applied Biosystems, Forster City, CA, USA) and analyzed on an Automated Capillary Electrophoresis Sequencer “3730 DNA Analyzer” (Applied Applied Biosystems, Forster City, CA, USA).

*Pseudo-nitzschia* sequences retrieved from GenBank for each marker (Suppl. Table 2) were aligned with sequences of *P. allochrona* using MAFFT ^33^, with the L-INS-i option. *Cylindrotheca fusiformis* and *Cylindrotheca* sp. were used as outgroup for 28S and *rbc*L, respectively, whereas *Fragilariopsis curta* and *F. cylindrus* were used as outgroups for 18S. No outgroups were included in the ITS alignment. Maximum likelihood (ML) was used for all markers. The substitution model used for each marker were selected through the Akaike information criterion (AIC) and the Bayesian information criterion (BIC) options implemented in MEGA X ^34^ respectively. ML analyses were performed in MEGA X and trees were built with 1,000 bootstrap replicates. All the phylogenetic trees were visualized using the interactive on-line tool iTOL (https://itol.embl.de, ^35^).

The analysis of the net evolutionary divergence, i.e. the number of base substitutions per site between ITS sequences of *P. allochrona* and of the most closely related species were conducted in MEGA XP. Standard error estimates were obtained through a bootstrapping procedure with 1,000 replicates.

The ITS2 secondary structure was predicted for sequences from strains SZN-B509 (Gulf of Naples) and SZN-B495 (Ionian Sea) for *P. allochrona* and NerD1 for *P. arenysensis* using RNA structure ^36^ with suboptimal structure parameters set as follows: maximum% energy difference 10, maximum number of structures 20, and window size 5. Format conversion (CT format to dot-bracket format) was performed with RNApdbee ^37^ and the 2D structures were drawn with VARNAv3.9 ^38^. The helices were labeled according to ^39^. Compensatory base changes (CBCs) detection was performed with 4SALE v1.7 ^40^ and hemi-compensatory base changes (H-CBCs) and other polymorphisms were observed manually.

### Mating experiments

Sexual reproduction experiments were conducted on 18 exponentially growing cultures of *P. allochrona* isolated in July 2016 (Suppl. Table 1). Prior to the experiments, the apical axis of 20 cells of each strain was measured in the light microscope. 153 couples of strains were mixed at concentrations of about 2,000 cells ml^−1^ each in six-well culture plates containing 4 mL of F/2 medium (Suppl. Table 3), which were incubated at the conditions described above. The mixed cultures were examined daily using a Zeiss Axiovert 200 light microscopes (Carl Zeiss, Oberkochen, Germany). The content of some wells where sexual reproduction was taking place was fixed and prepared for SEM observations. By convention, female sex was attributed to the strain to which zigotes were attached as recognised in crosses between strains with cells of different sizes, and to all strains sexually incompatible with those ones.

Sexual compatibility was tested by crossing two strains of opposite mating type (BB16 and CM63) of *P. arenysensis* between them and with each of 4 strains of *P. allochrona* (8B5, 8C3, 8C5 and 9A2) as described above.

### Ecological analysis

Cell densities of *P. delicatissima*-like morphotypes were assessed in 1,154 phytoplankton samples collected by Niskin bottles at surface at the LTER-MC stations beweekly from January 1984 through July 1991 and weekly from February 1995 to December 2015. Samples fixed with buffered 37% formaldehyde (1.6% final concentration) were examined and counted following the Utermöhl method ^41^ with a Zeiss Axiovert 200 light microscope (LM) (Carl Zeiss, Germany). Simultaneous data for environmental variables (temperature, salinity and nutrients) were collected and quality controlled as described in ^42^. *P. delicatissima*-like morphs occurring in the second half of the year were arbitrarily assigned to *P. allochrona*, based on their recurring seasonal pattern (see results). Those recorded in the first part of the years were assigned to other *P. delicatissima*-like species, which included *P. delicatissima* sensu stricto, *P. arenysensis* and *P. dolorosa*. The environmental niches of *P. allochrona* and the other *P. delicatissima*-like morphs were explored with the co-inertia analysis Outlying Mean Index (OMI) ^43^, which generates ordination axes that maximize the separation between species occurrences in a multivariate environmental space. The positions of the species in the environmental space, representing the deviation of the species from a theoretical ubiquitous species occurring under all environmental conditions, were compared to simulated values (1,000 random permutations) under the null hypothesis that each species is uninfluenced by the environment. The environmental map was defined based on surface values of physical (temperature and salinity) and chemical parameters (dissolved inorganic nitrogen, silicates and phosphates) and by photoperiod.

## Results

### Diagnosis

*Pseudo-nitzschia allochrona* Zingone, Percopo et Sarno sp. nov. (Fig. 1, A–J; Suppl. Table 3)

**Figure 1.**
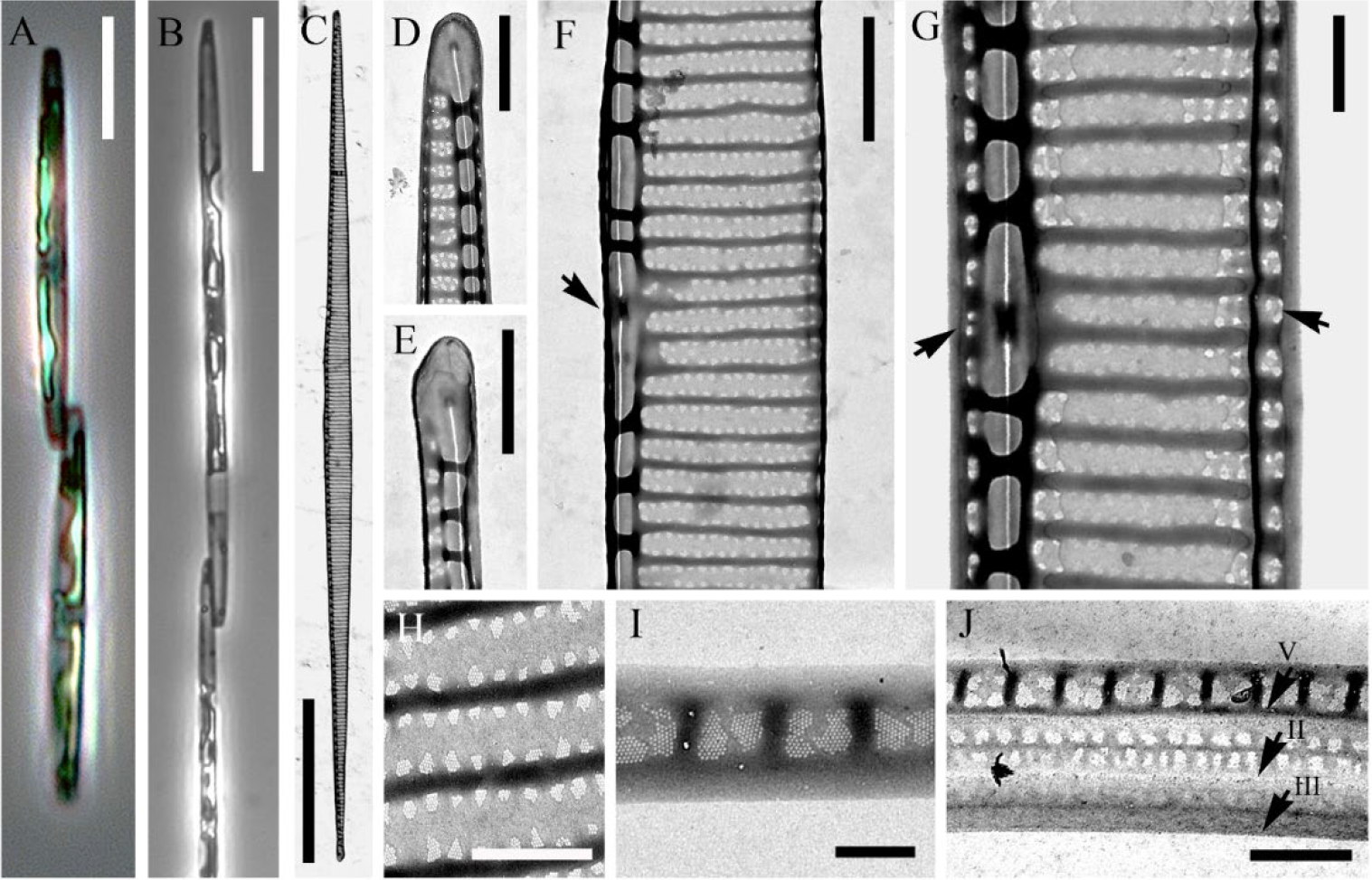
Morphology of *Pseudo-nitzschia allochrona* sp. nov.. LM (A, B) and TEM micrographs (C-J). (A) Cells in girdle view, strain MC784 4 II. (B) Long cells formed following sexual reproduction (cross 9A2×9C3A, Suppl. Table 3), girdle view. (C) Whole valve. (D) Valve end. (E) Valve end. (F) Central part of the valve face with central nodule. (G) Central part of the valve with central nodule and mantle (arrowheads). (H) Detail of the valve striae with two rows of pores typical of the *P. delicatissima-*complex. (I) Detail of the valvocopula. (J) The three cingular bands, with arrows indicating their borders: V = valvocopula, II = second cingular band, III = third cingular band. (A): strain MC784 4 II, (B): (C-J): strain B109. Scale bars: (A) = 5 μm, (B)= 20μm, (C) = 10 μm, (D, E, F)= 1 μm, (G, H, J) = 0.5 μm, (I)= 0.2 μm..

Cells lanceolate with pointed ends forming stepped colonies. Apical axis 22-84 μm, transapical axis: 1.4-2.1 μm. Valves with larger central interspace, striae with two rows of irregular poroids, 10-12 in 1 μm. 20-26 fibulae and 34-44 interstriae in 10 μm. Cingulum with three open bands: i) valvocopula, 1-2 poroids high and 2 poroids wide, with 46-50 striae in 10 μm, ii) second band, with a longitudinal silicified line flanked by two rows of poroids, and iii) third band almost unperforated.

*Holotype*: Slide of strain SZN-B501 deposited at the Museum of the Stazione Zoologica Anton Dohrn (SZN).

*Isotype*: Fixed material of SZN-B501 deposited at the Museum of the SZN, registered as XXXX.

*Epitype*: Molecular characterization: DNA sequences for rDNA LSU, ITS and *rbc*L of strain SZN-B501 are deposited in GenBank with accession numbers XXXX, XXXX and XXXX, respectively.

*Type locality*: LTER-MareChiara, 40° 48’50’’ N; 14° 15’ 0’’ E, Gulf of Naples (Mediterranean Sea).

*Etymology*: the epithet (allos: other, chronos: time) refers to the distinct phenology of the species compared to other *Pseudo-nitzschia delicatissima*-like species from the type locality.

In light microscopy *P. allochrona* shows the typical features of *P. delicatissima*-like species, including thin valves, moderate length and relative thickness of cell ends in lateral view, resulting in a pronounced step in cell chains (Fig. 1A, B). No morphological, morphometric or ultrastructural characters investigated (Fig. 1 C-J) discriminate *P. allochrona* from several *P. delicatissima*-like species, including the ones found in the Gulf of Naples, namely, *P. delicatissima*, *P.arenysensis* and *P. dolorosa* (Suppl. Table 3).

The size of 18 strains of *P. allochrona* tested in breeding experiments ranged between 22 and 54 μm (38.5 ± 4.2 μm; n = 364) in apical axis. All strains mated in numerous successful crosses that allowed to identify two groups of 10 and 8 strains of opposite mating type (Suppl. Table 4). Gametangia of different strains and at times of different size paired about one day after the inoculum (Fig. 2A). Couples of zygotes, initially spherical (Fig. 2B), modified synchronously into elongate auxospores (Fig. 2C, D) attached to the frustule of one empty gametangium (the ‘female’ strain by convention). Mature auxospores were often curved, with a clear bulge in the center, a cap at each end and a transversal perizonium with fairly ornamented bands (Fig. 2E). Initial cells (apical axis 72–84 μm, 79.9 ±2.3 μm; n = 21) were also slightly curved (Fig. 2 F) but regained the straight shape after the first vegetative divisions (Fig. 1B).

**Figure 2.**
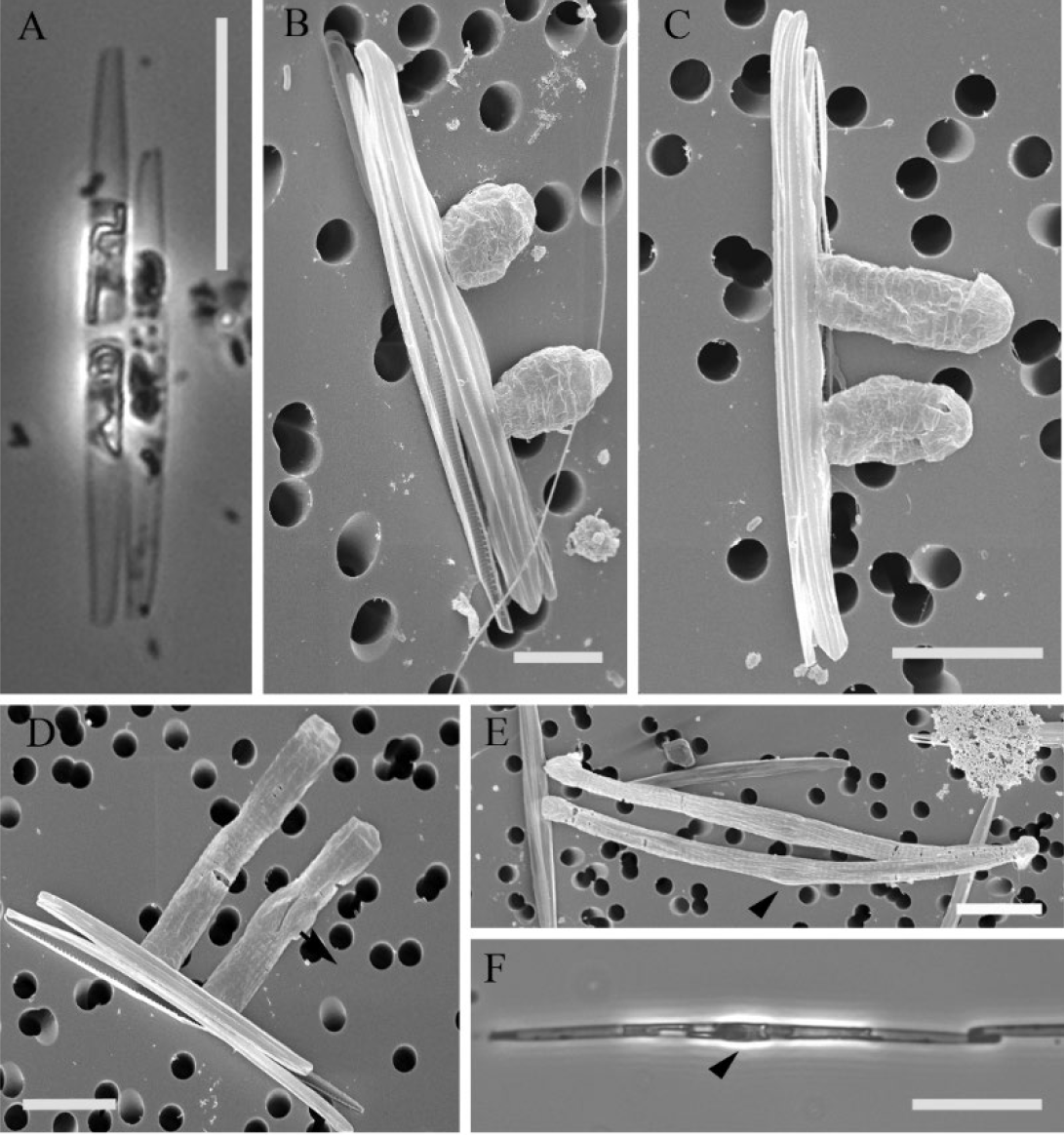
Life stage of *Pseudo-nitzschia allochrona* sp. nov. during sexual reproduction. LM (A, F) and SEM (B-E). (A) Two paired gametangia of opposite mating types and different sizes, cross of strains 9A2×9C5. (B) Gametangia with two zygotes connected to the parental valve, cross 9B4×9C5. (C) Early auxospores, cross 9B4×9C5. (D) Elongated auxospores, cross 9B4×9C3a. (E) Mature auxospores with a bulge in the center (arrowheads), still connected to the parental valve, cross 9B4×9C5. (F) Long cell following the first divisions with the central bulge visible (arrowhead), cross 9A2×9C5. Scale bars: (A and F) = 20 μm, (B)= 5μm, (C-E) = 10 μm.

All 4 markers investigated (18S, ITS, 28S, and *rbc*L) showed *P. allochrona* as distinct from all known congeneric species and clustering with moderate to high support with other *P. delicatissima*-like species (Fig. 3, Suppl. Fig.1). The new species was closely related to *P. arenysensis* in all phylogenies but the 18S one, where it was associated to *P. delicatissima* in a poorly supported clade The net evolutionary divergence in ITS between *P. allochrona* and *P. arenysensis* (0.042 ±0.010) was lower than that with *P. delicatissima* (0.067±0.011) (Suppl. Table 5).

**Figure 3.**
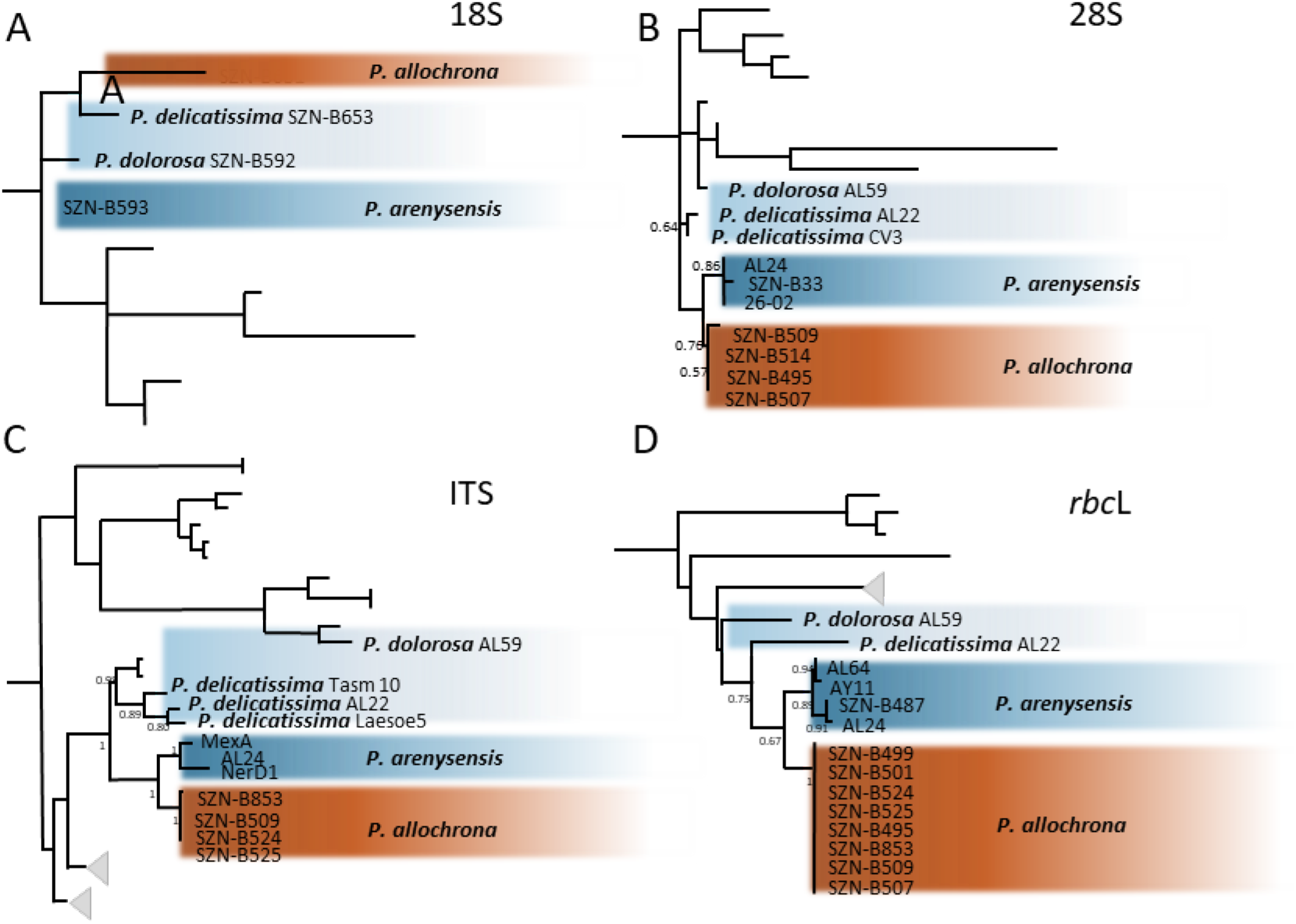
Maximum Likelihood phylogenies of the *P. delicatissima*-complex species living in sympatry in the Gulf of Naples. Excerpts from Suppl. Fig. 1 presenting the complete phylogenetic trees. A) 18S; B) 28S; C) ITS; and D) *rbc*L. The new species *P. allochrona* is well separated from the closely related species *P. delicatissima, P. dolorosa and P. arenysensis* in the four markers, and sister to *P. arenysensis* in all supported phylogenies, being closer to *P. delicatissima* only in the non-supported 18S phylogeny.

The ITS2 secondary structure of *P. allochrona* showed four main helices (Helix I-IV) and one pseudo-helix (IIa), with a pyrimidine–pyrimidine mismatch in helix II, similar to other congeneric species ^25^ (Fig. 4). Compared to its closest relative *P. arenysensis*, *P.allochrona* ITS2 had one CBC in helix I (C-G in *P.allochrona* ↔ G-T in *P. arenysensis*) and two hemi-CBCs in helices I (C-G ↔ G-T) and III (G-T↔ AT). In helix I, two deletions (AGTGT and ATTCT) determined the loss of one internal loop pyrimidine-pyrimidine and a change in the hairpin loop structure. A single SNP was found in the internal loop (C→T) of helix II and four SNIPs in the internal loops (TTT →CAC and A→C) of helix III, where a deletion of GG produced the loss of an internal pyrimidine-pyrimidine internal loop. The stem length of helix IV was shorter because of three deletions (GGTT, ATAG and ATTGTAC), which also determined changes in the conformation of the hairpin loop.

**Figure 4:**
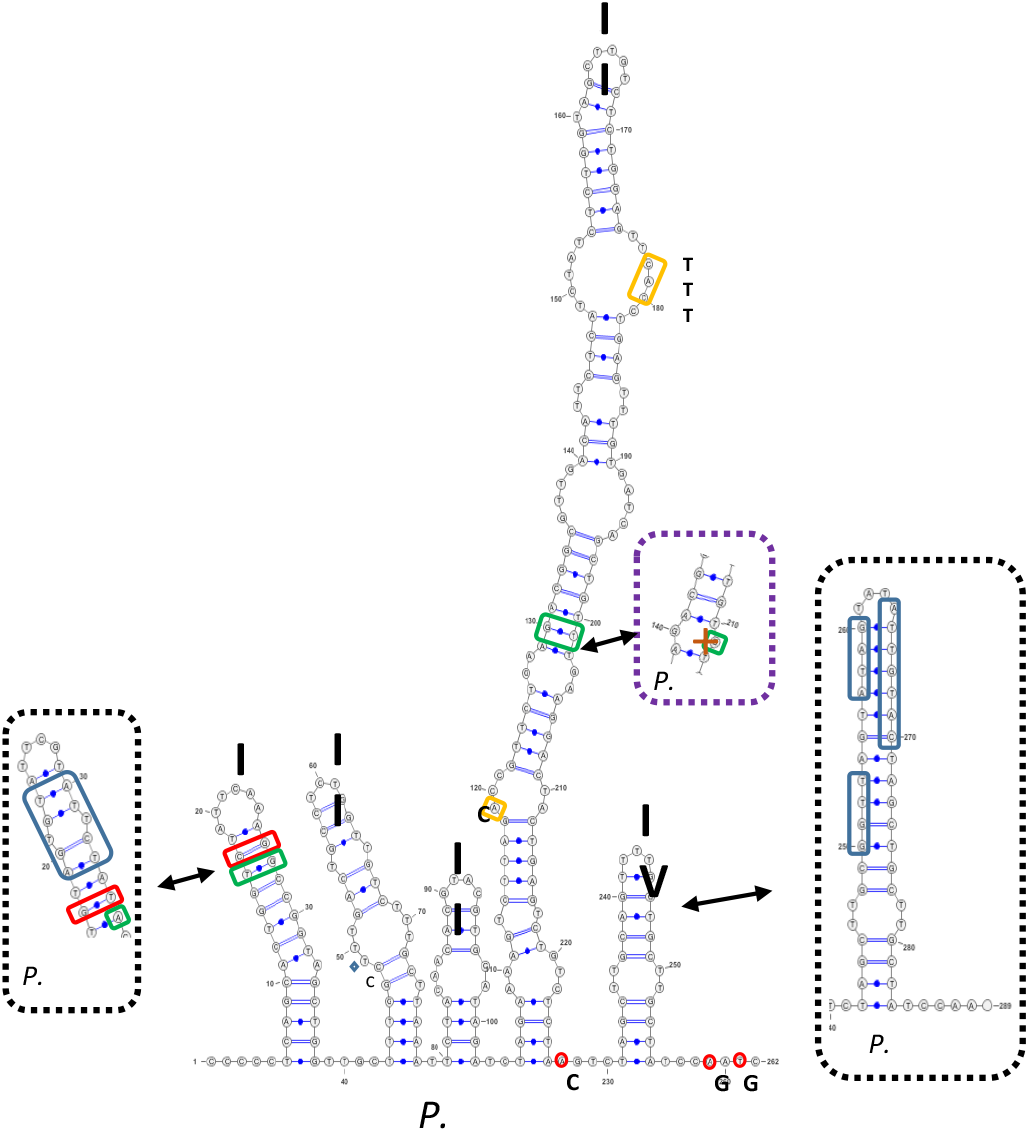
ITS2 secondary structure of *P. allochrona* (SZN-B509) and comparison to that of the sibling species *P. arenysensis* (NerD1). Dashed black boxes: indels in the alignment of the two species; red boxes: CBCs; green boxes: Hemi-CBCs; orange boxes: SNPs; and dashed purple box: an Hemi-CBC generating an internal pyrimidine-pyrimidine loop **(+**). Red circles reflect ITS intra-specific polymorphisms between *P. allochrona* strains from the Gulf of Naples and those from the Ionian Sea (SZN-B495).

Two interfertile *P. arenysensis* strains crossed with four *P. allochrona* strains of opposite mating type did not show any sign of sexual reproduction between the two species (Suppl. Table 4).

Toxin analyses of strains SZN-B524 and SZN-B495 did not reveal the presence of domoic acid.

At the LTER-MC station, *P. delicatissima*-like species showed a first bloom period from March through May, and a second one from late June through mid-September (Fig. 5 A, B), with minima generally in late spring and late autumn-winter but occasional peaks also in the latter periods. Through molecular analyses, all 62 *P. delicatissima*-like strains isolated from the Gulf of Naples from late June onwards over 10 years were attributed to *P. allochrona*, while 187 strains isolated in the first part of the year over multiple years all belonged to *P. arenysensis* and *P. delicatissima*, and more rarely to *P. dolorosa* (Fig. 5C). Based on this clear temporal separation, also supported by metabarcoding results over different years (see discussion), we arbitrarily assigned records of summer-autumn *P. delicatissima*-like morphotypes in the LTER-MC time series to *P. allochrona* and winter-spring ones to the remainders species in order to assess the specificity of the ecological niche of the new species. The first OMI axis of the niche analysis of all *P. delicatissima*-morphs (Fig. 5D) accounted for most of the variability (75.32%) and described the environmental gradient from early-spring conditions, with relatively high nutrient levels and low temperature, to higher temperature and poorer nutrient concentrations in summer. The second OMI axis (27.68%) defined the environmental gradient of light and salinity separating the average habitat conditions occurring in late-autumn and late-spring. *P. delicatissima* morphs’ niche positions significantly deviated from the origin of the multivariate space (p-value < 0.001) pointing at the relationship of their seasonal distribution with environmental conditions. The clear separation along the first OMI axis reflected the association of *P. allochrona* with higher temperature in a relatively nutrient-poor environment compared to other *P. delicatissima*-like morphs. In either conditions, longer days appeared to favour more numerous and intense blooms (Fig. 5C, D). Over more than 30 years of sampling at the LTER-MC stations, annual densities of *P. delicatissima*-morphs occurring in the second part of the year, here attributed to *P. allochrona,* were null or very low in the first years of the time series (1984-1989), while their abundance increased particularly over the last 15 years, in some years overtaking that of the other species (Fig. 5E).

**Figure 5.**
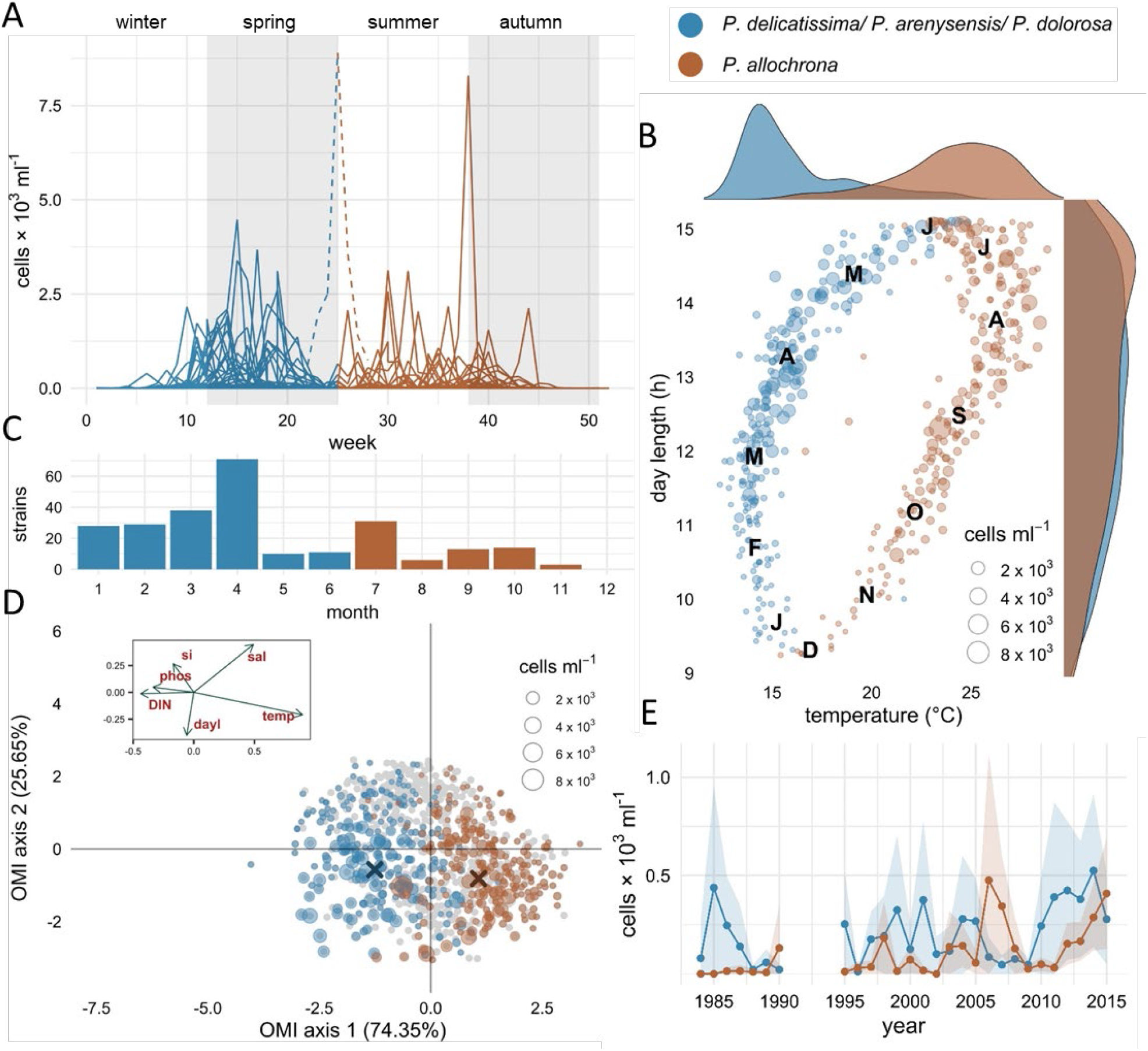
Distribution and ecology of species of the *P. delicatissima*-complex in the Gulf of Naples. A) Annual distribution (1984-2015) of *P. allochrona* and of the other *P. delicatissima*-like species (*P. delicatissima, P. arenysensis* and *P. dolorosa*, lumped) from light microscopy observations. *P. allochrona* is distinguished from the others based on its recurring occurrence in summer-autumn; lines represent different years. The exceptional peak in late June-early July 2014 (dashed line), not attributable to either species, was not included in the niche analysis of panel D. B) Separation of *P. delicatissima*-like species in the seasonal space identified by day length and temperature values. Letters are month names’ initials. C) Annual distribution of *P. allochrona* and the three other *P.delicatissima-*like species based on the isolation date of numerous strains (62 and 187, respectively) identified with molecular methods over the years 2004-2016. D) Ecological niche analysis of *P. allochrona* and the three other *P. delicatissima*-like species, showing *P. allochrona* separated from that of the other species along the OMI axis 1, highly correlated with temperature (temp) and negatively correlated with nutrients (DIN: dissolved inorganic nutrients; phos: phosphate; si: silicate). More and larger dots in the 3^rd^ and 4^th^ quadrants indicate higher frequency and abundance for all species in periods of greater daylength (dayl) and lower salinity (sal). Grey dots represent samples with no species of the *P. delicatissima-*complex. E) Interannual density variations (lines: annual average values; shadowed areas: C.I. 95%) of the *P. delicatissima-*like species; *P. allochrona* was hardly detected over the first years of the time series.

## DISCUSSION

Several different evidences in this study support *P. allochrona* as a new cryptic species within the *P. delicatissima-*complex. Morphologically undistinguishable by definition, its individuality is clearly seen in the molecular signature of three nuclear and one chloroplastic sequences that are commonly used for species delimitation in diatoms. Further, marked differences and conformational changes in the ITS secondary structure compared to its closest relative *P. arenysensis* and the failure to perform sexual reproduction indicate mating incompatibility between the two sister species ^25,44^.

Nevertheless, it is the phenological signature the most conspicuous character that distinguishes *P. allochrona* from the other *P. delicatissima*-like species living in sympatry in the Gulf of Naples. Since its first record in 2004 (as ‘*Pseudo-nitzschia* new genotype’) in a clone library-based DNA-metabarcoding study of the 28S rDNA fragment ^45^, *P. allochrona* has always been the only species of the group found in summer-autumn, never showing up among the more than 187 strains of the *P. delicatissima*-complex retrieved in winter-spring in the area over more than 10 years, all invariably identified as *P. delicatissima*, *P. arenysensis* or *P. dolorosa*. The temporal segregation of these species is also confirmed by a seasonal overview based on 28S rDNA clone-library, in which *P. allochrona* (as *P. delicatissima* IV) was responsible for the late summer-early autumn blooms of 2009 and 2010 ^18^, while a recent three-ys HTS study based on 17,763 environmental 18S rDNA-V4 barcodes has again detected *P. allochrona* in late summer in 2011 and 2013 ^46^. In all the three metabarcoding studies, *P. delicatissima/P.dolorosa* (not separated by the V4-18SrDNA marker) and *P. arenysensis* overlap to a large extent in their occurrences, being the main contributors to the spring peak of this species-complex. Peaks of *P. arenysensis* in some years followed and in other years preceded those of *P. delicatissima,* which actually co-occurred with *P. allochrona* in late June-early July of some years.

The phenological segregation between *P. allochrona* and its cryptic congeneric species matches the differences in the environmental conditions of different periods of the year, whereby temperature mainly and nutrient levels to some extent seem to be the drivers of the separation of the two annual peaks of the species complex. While nutrients should not be limiting in either seasonal context ^23^, the differences of ca. 10 °C between spring and summer-autumn temperature values suggests eco-physiological variations between the species responsible for blooms at different times of the year. In *Thalassiosira rotula*, changes in the genetic structure among populations also correlate strongly with water temperature at the spatial and temporal scale ^11^, whereas a salinity gradient drove spatial patterns of genetic diversity in the case of *Skeletonema marinoi* in the Baltic Sea ^47^. The relationship between neutral and functional genetic diversity within and among species is complex ^48^, but the adaptive response under new environmental condition can be very rapid in diatoms ^49,50^, while profound functional genomic differences have been found between sister species in the prasinophyte *Ostreococcus* ^51^. In the case of *P. allochrona*, biochemical differences between this species (as *Pseudo-nitzschia* cf. *delicatissima*) and its sister species *P. arenysensis* and *P. delicatissima* have been found in lipoxygenase enzymes mediating the metabolism of eicosapentaenoic acid ^19^, which provides a first indication of functional differences within this group of species.

Morphological identity and relatively small genetic distance between *P. arenysensis* and *P. allochrona* point at a possibly recent speciation event. Despite numerous recent studies focusing on the genus *Pseudo-nitzschia* all over the world, so far *P. allochrona* has only been found in the Gulf of Naples ^18,19,45^, in the Ionian Sea (this paper) and more recently in the Adriatic Sea ^52–54^. In the latter area *P. allochrona* (as *P.* cf. *arenysensis*) has only been found in summer-autumn, like in the Gulf of Naples and Ionian Sea, while *P. arenysensis* has never been detected. Although speciation by geographic isolation cannot be ruled out, it is tempting to postulate that the separation between the two sister species, *P. allochrona* and *P. arenysensis* may have occurred in sympatry in the Gulf of Naples. Habitat heterogeneity promotes species diversity at both the ecological and evolutionary time scale, preventing competitive exclusion and providing new niches to be occupied by different, co-existing species, eventually leading to ecological speciation ^7,55^. Whereas spatial partitioning of ecological conditions is hard to occur in the planktonic realm, especially for phototrophs, environmental factors can vary considerably along the year in strongly seasonal environments, such as the Mediterranean Sea, whereby time can replace space in creating the habitat heterogeneity required for ecological divergence, thus leading to allochronic speciation. A similar example of divergence by time in the same genus is offered by *Pseudo-nitzschia galaxiae*, which blooms in the Gulf of Naples in three different periods with populations of three size classes ^56^. The size classes actually correspond to distinct ribotypes ^57^ that are also retrieved as distinct in e-DNA metabarcoding studies ^18^. Whether the three *P. galaxiae* populations are separate species needs further investigation, but the coherent pattern observed in size ranges, genetics and timing of the blooms provides a further possible case of isolation by time and allochronic speciation processes.

Genetic divergence among populations occurring at different times has been observed in various phytoplankton groups ^11,58–61^, suggesting that temporal segregation could be a general mechanism for speciation in marine protists. Yet phenology has rarely been considered a stable, endogenous character in phytoplankton, whereby the alternation of different species over the seasons is postulated to be strictly driven by changes in environmental conditions. Nevertheless, annually recurrent patterns are seen in long term trends of nano- and microphytoplankton species of the Gulf of Naples ^23,62^, while recent massive eDNA sequencing also revealed recurrent occurrence of picoplanktonic taxa against marked environmental variability ^63,64^, which can be explained based on endogenous rhythms and stability of phenological characteristics.

Population dynamics in *Pseudo-nitzschia* species is compatible with a scenario of stable phenological rhythms, as a synchronous growth timing maximises encounter probability, which is a key element for sexual reproduction in heterotallic species in a highly dispersive habitat. Sexual reproduction in natural *Pseudo-nitzschia* populations has been inferred from shifts in cell size ^65^ and rarely observed, but interestingly in September 2008 a massive sexual event was recorded at station LTER-MC at the end of a bloom of a *P. delicatissima*-like species, which was probably *P. allochrona* based on the time of the year ^66^. As *Pseudo-nitzschia* species are not known to produce benthic stages, sparse cells persist in the plankton outside the density peak time and occasionally may give rise to blooms in other periods of the year. In this context, slight variations in peak times and unusual blooms in some years, as the one recorded in June 2015, suggest that the attempt to colonize new time windows may occur frequently in this genus or in phytoplankton at large, which could be seen as wandering across the seasons in search of new ecological niches.

The mechanisms underlying allochronic speciation, and speciation in general, are not easy to clarify ^48^, as both neutral and selective process could be involved. Time and environmental variables covary, making it difficult to discern their respective contribution to the patterns observed in this study. One possibility is that ecological segregation arises as a consequence of phenological shifts in the timing of the maximum abundance, a mechanism that would reduce competition for resources among individuals within a population ^67^. While most individuals respond rapidly and similarly to the relevant environmental cues, phenological characters often follow a skewed distribution ^68^, the long tail implying that a small part of the population may experience new environmental conditions during key life-cycle steps, such as reproduction. In such conditions that promote assortative mating, reduced gene flow can eventually lead to reproductive isolation between groups ^13^. This situation is somewhat analogous to a founder effect, where a subsample of the population colonizes new niches, diverging from the mother population via genetic drift ^69^. Variations of the phenological timing could be favoured by the phenotypic plasticity that is typical of microalgae and/or rapid adaptation to new ecological conditions ^49^, or by selective processes acting on standing infraspecific diversity ^48^, which is also quite wide in phytoplankton populations ^70^.

The separation by time could be rapid enough as to be observable on decadal time scales, as no clearcut boundary exists between the time-scales of ecological and evolutionary processes ^71^. Contemporary evolution could occur especially in microbial organisms, which are characterized by high growth rates and a few to several tens of generations over a single bloom season. In fact, rapid genetic variations take place in phytoplankton ^70,72^ and interannual variations in the genetic structure of the population were actually observed in another species of the genus, *P. multistriata* ^73^, which also showed intermittent periods of weak and strong intraspecific diversity over the same bloom season ^74^.

Not different from terrestrial plants, unicellular aquatic phototrophs can capture information from light and possess genes that are involved in the regulation of biological rhythms at various scales ^75,76^. Circa-annual rhythms and regular phenological patterns associated to photoperiodic response suggest that these microorganisms may also be able to measure the time of the year ^63,77^. In this perspective, diversity in biological rhythms of plankton would be an optimal substrate for their evolutionary changes, contributing to isolation by time and speciation. In support of this proposition, future studies should address genetic variations in sympatric populations and sibling species over the seasonal and interannual time-scale and assess their relationships with functional adaptation through the analysis of neutral and adaptive genetic variations, an approach now made possible by the accessibility of metagenomics and metatranscriptomic technologies. In this respect, the present study highlights the potential of combining molecular and ecological information over long term time series in order to trace the eco-evolutionary dynamics and shed light on speciation mechanisms in the plankton.

## Abbreviations

AIC: Akaike information criterion
BI: Bayesian Inference
BIC: Bayesian information criterion
CBC: compensatory base change
DA: domoic acid
HCBC: hemi-compensatory base change
LTER: Long Term Ecological Research
ML: Maximum Likelihood

## Acknowledgments

The authors thank A. Passarelli, F. Tramontano, M. Cannavacciuolo, and G. Zazo for sampling and all LTER-MC team and the crew of the R/V Vettoria for assistance during the work at sea, C. Minucci for help with culture isolation and maintenance and molecular characterization of the strains, and R. Graziano and F. Iamunno for electron microscopy assistance. Physical and chemical data from the station LTER-MC were kindly provided by F. Margiotta (SZN).

## Author contribution

AZ and DS conceived and organised the present research. IP, MVR and RP provided and analysed the molecular data, DS, AZ and IP provided and analysed the morphological data, LL conducted the statistical ecological analyses and RR the biochemical analyses. AZ, IP and MVR drafted the manuscript, with contributes from all other authors who read and approved the final version. AZ coordinated the research team.

