## Supplemental Figure 1 for "Phenological segregation suggests speciation by time in the planktonic diatom *Pseudo-nitzschia allochrona* sp. nov."

Fig. S1: Maximum-likelihood phylogenies of the genus *Pseudo-nitzschia*. Squares indicate the part of the trees shown in Fig. 1. A) 18S; B) 28S; C) ITS; D) *rbc*L. Only bootstrap values > 0.5 are shown.

A

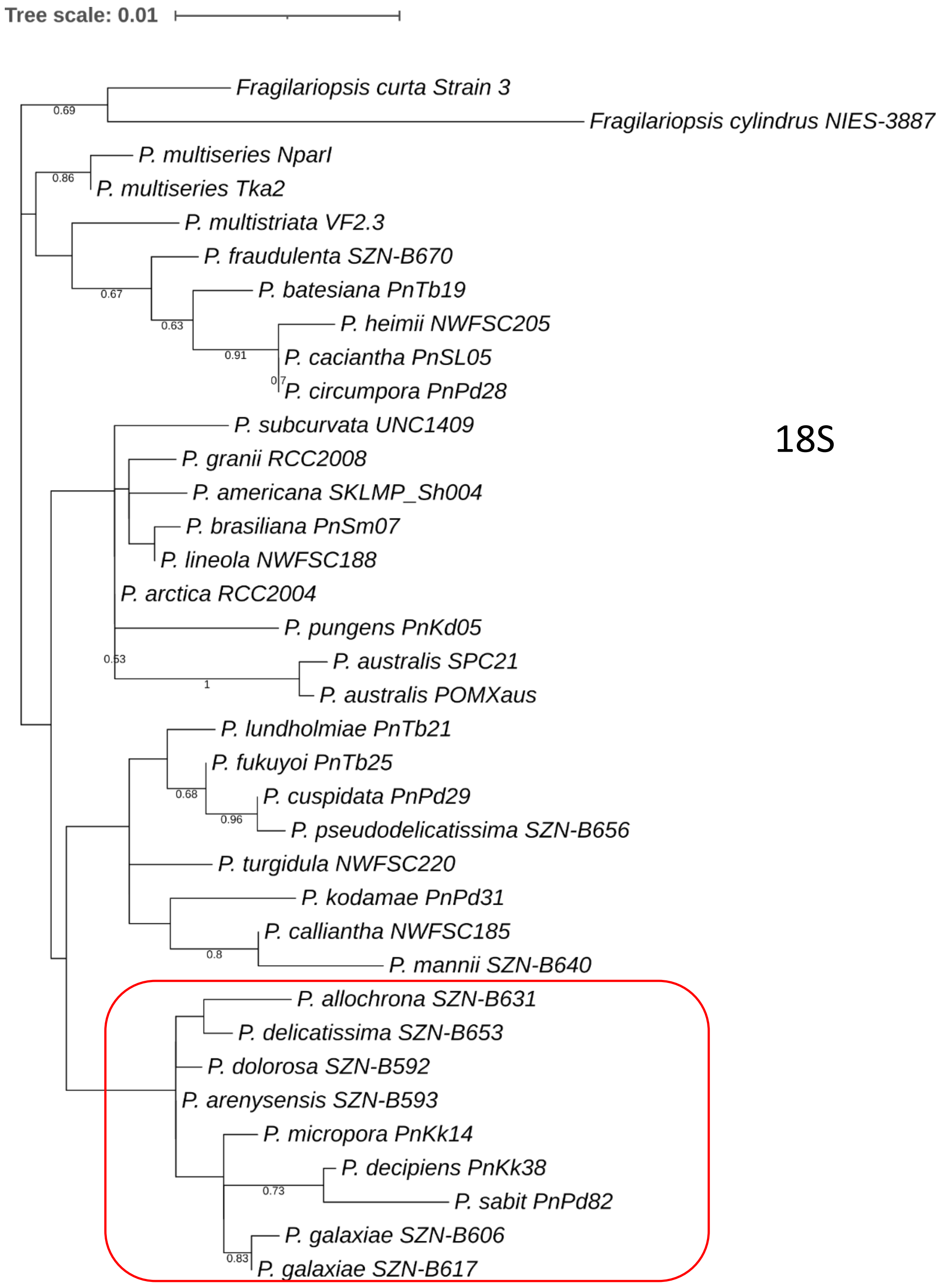

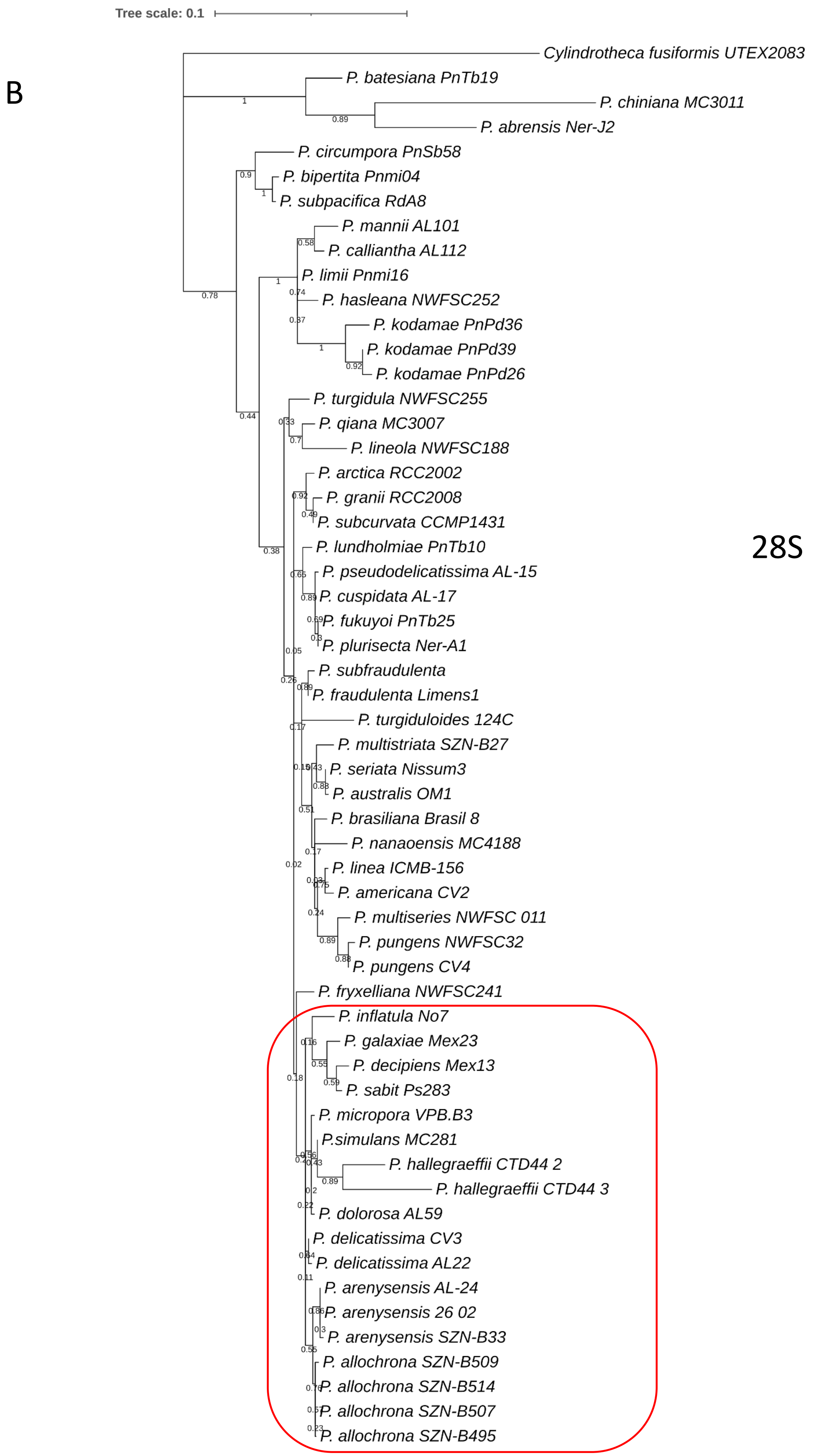
Tree scale: 0.1
B
28S
Cylindrotheca fusiformis UTEX2083
P. batesiana PnTb19
P. chiniana MC3011
P. abrensis Ner-J2
P. circumpora PnSb58
P. bipertita Pnmi04
P. subpacifica RdA8
P. mannii AL101
P. calliantha AL112
P. limii Pnmi16
P. hasleana NWFSC252
P. kodamae PnPd36
P. kodamae PnPd39
P. kodamae PnPd26
P. turgidula NWFSC255
P. qiana MC3007
P. lineola NWFSC188
P. arctica RCC2002
P. granii RCC2008
P. subcurvata CCMP1431
P. lundholmiae PnTb10
P. pseudodelicatissima AL-15
P. cuspidata AL-17
P. fukuyoi PnTb25
P. plurisecta Ner-A1
P. subfraudulenta
P. fraudulenta Limens1
P. turgiduloides 124C
P. multistriata SZN-B27
P. seriata Nissum3
P. australis OM1
P. brasiliana Brasil 8
P. nanaoensis MC4188
P. linea ICMB-156
P. americana CV2
P. multiseries NWFSC 011
P. pungens NWFSC32
P. pungens CV4
P. fryxelliana NWFSC241
P. inflatula No7
P. galaxiae Mex23
P. decipiens Mex13
P. sabit Ps283
P. micropora VPB.B3
P.simulans MC281
P. hallegraeffii CTD44 2
P. hallegraeffii CTD44 3
P. dolorosa AL59
P. delicatissima CV3
P. delicatissima AL22
P. arenysensis AL-24
P. arenysensis 26 02
P. arenysensis SZN-B33
P. allochrona SZN-B509
P. allochrona SZN-B514
P. allochrona SZN-B507
P. allochrona SZN-B495

C

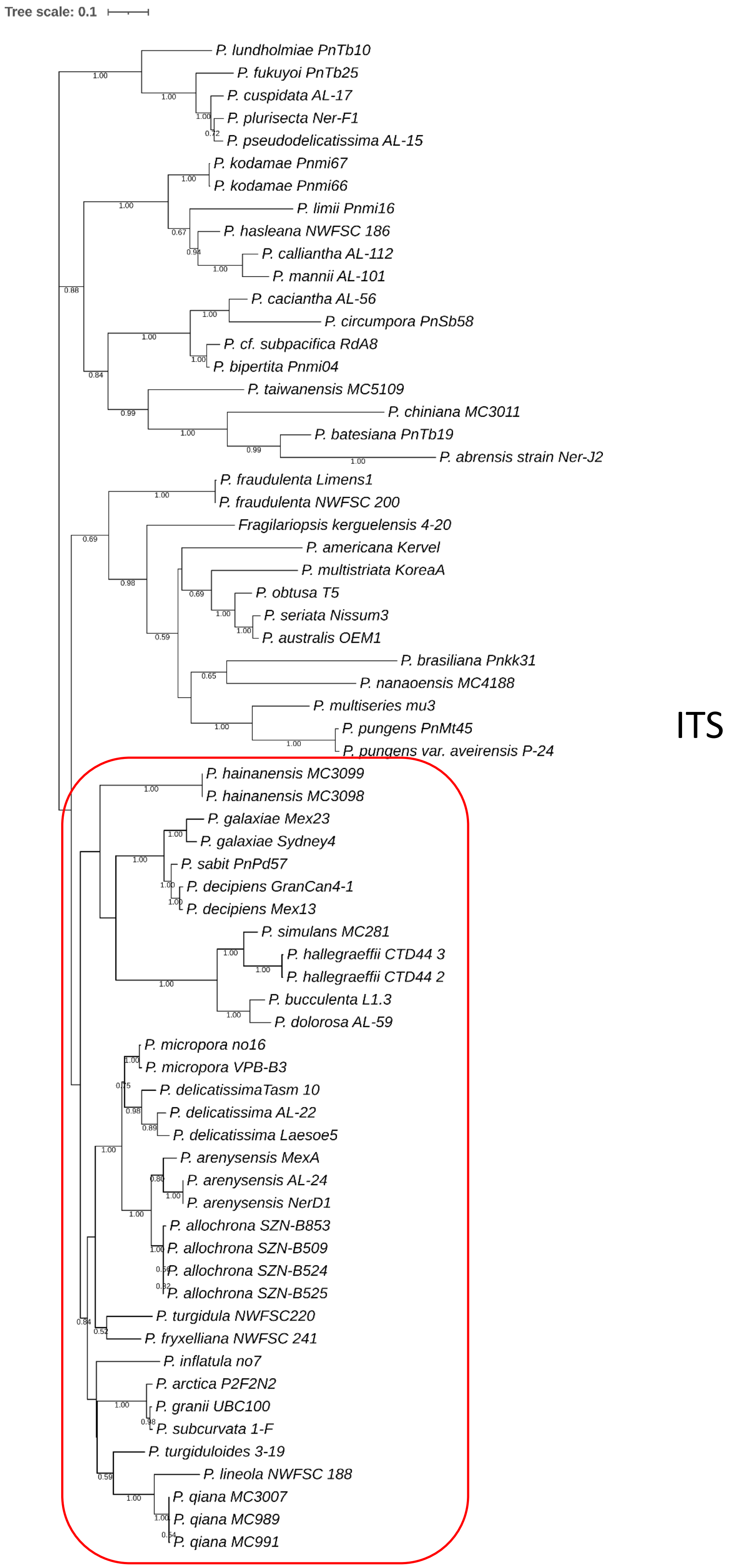

Tree scale: 0.1
P. lundholmiae PnTb10
P. fukuyoi PnTb25
P. cuspidata AL-17
P. plurisecta Ner-F1
P. pseudodelicatissima AL-15
P. kodamae Pnmi67
P. kodamae Pnmi66
P. limii Pnmi16
P. hasleana NWFSC 186
P. calliantha AL-112
P. mannii AL-101
P. caciantha AL-56
P. circumpora PnSb58
P. cf. subpacifica RdA8
P. bipertita Pnmi04
P. taiwanensis MC5109
P. chiniana MC3011
P. batesiana PnTb19
P. abrensis strain Ner-J2
P. fraudulenta Limens1
P. fraudulenta NWFSC 200
Fragilariopsis kerguelensis 4-20
P. americana Kervel
P. multistriata KoreaA
P. obtusa T5
P. seriata Nissum3
P. australis OEM1
P. brasiliana Pnkk31
P. nanaoensis MC4188
P. multiseries mu3
P. pungens PnMt45
P. pungens var. aveirensis P-24
P. hainanensis MC3099
P. hainanensis MC3098
P. galaxiae Mex23
P. galaxiae Sydney4
P. sabit PnPd57
P. decipiens GranCan4-1
P. decipiens Mex13
P. simulans MC281
P. hallegraeffii CTD44 3
P. hallegraeffii CTD44 2
P. bucculenta L1.3
P. dolorosa AL-59
P. micropora no16
P. micropora VPB-B3
P. delicatissimaTasm 10
P. delicatissima AL-22
P. delicatissima Laesoe5
P. arenysensis MexA
P. arenysensis AL-24
P. arenysensis NerD1
P. allochrona SZN-B853
P. allochrona SZN-B509
P. allochrona SZN-B524
P. allochrona SZN-B525
P. turgidula NWFSC220
P. fryxelliana NWFSC 241
P. inflatula no7
P. arctica P2F2N2
P. granii UBC100
P. subcurvata 1-F
P. turgiduloides 3-19
P. lineola NWFSC 188
P. qiana MC3007
P. qiana MC989
P. qiana MC991
ITS

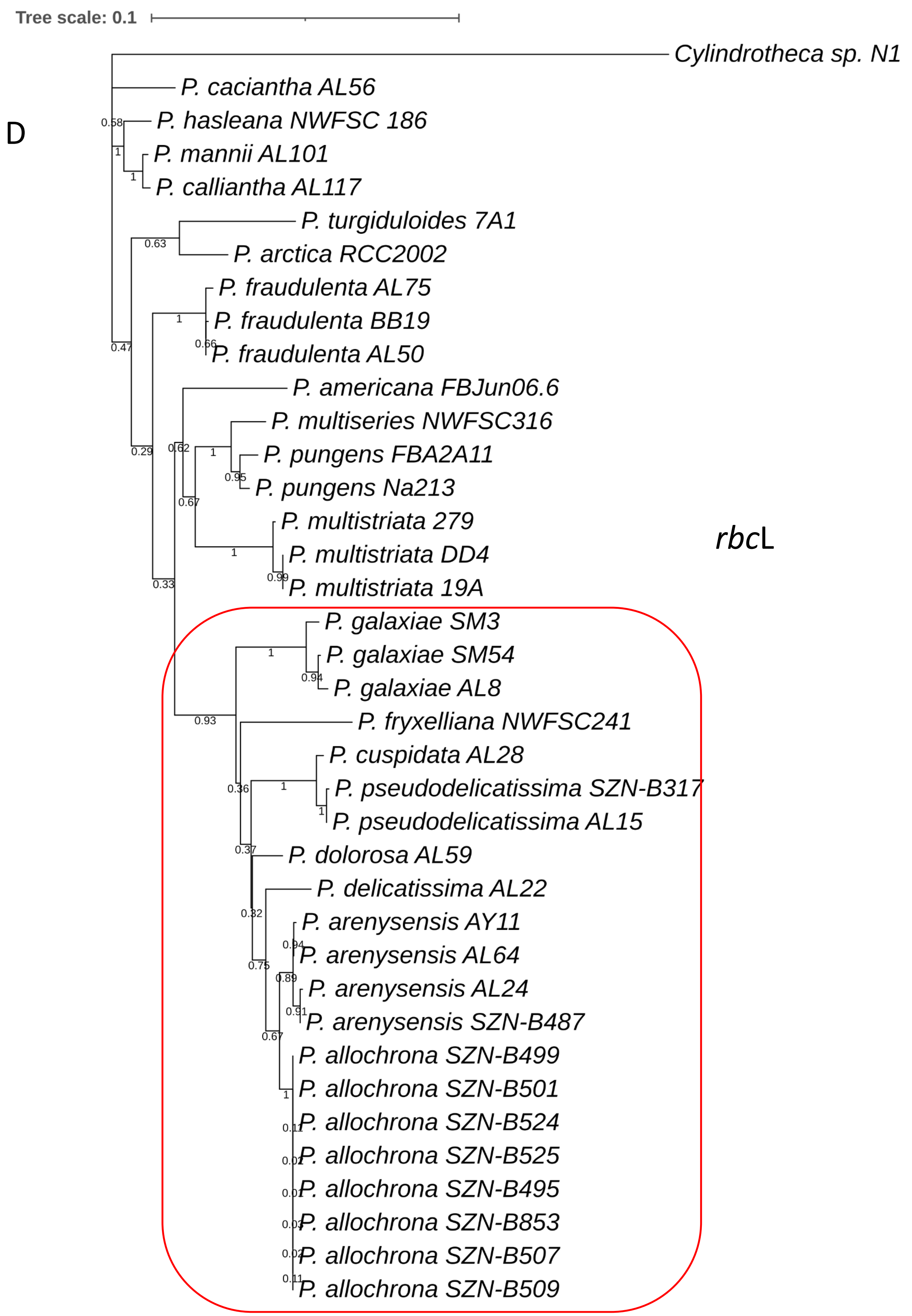

Tree scale: 0.1
D
rbcL
Cylindrotheca sp. N1
P. caciantha AL56
P. hasleana NWFSC 186
P. mannii AL101
P. calliantha AL117
P. turgiduloides 7A1
P. arctica RCC2002
P. fraudulenta AL75
P. fraudulenta BB19
P. fraudulenta AL50
P. americana FBJun06.6
P. multiseries NWFSC316
P. pungens FBA2A11
P. pungens Na213
P. multistriata 279
P. multistriata DD4
P. multistriata 19A
P. galaxiae SM3
P. galaxiae SM54
P. galaxiae AL8
P. fryxelliana NWFSC241
P. cuspidata AL28
P. pseudodelicatissima SZN-B317
P. pseudodelicatissima AL15
P. dolorosa AL59
P. delicatissima AL22
P. arenysensis AY11
P. arenysensis AL64
P. arenysensis AL24
P. arenysensis SZN-B487
P. allochrona SZN-B499
P. allochrona SZN-B501
P. allochrona SZN-B524
P. allochrona SZN-B525
P. allochrona SZN-B495
P. allochrona SZN-B853
P. allochrona SZN-B507
P. allochrona SZN-B509
