## supplemental Tables 1-5 for "Phenological segregation suggests speciation by time in the planktonic diatom *Pseudo-nitzschia allochrona* sp. nov."

**Table S1: List of strains of *Pseudo-nitzschia allochrona* used in the present study for electron microscopy in (EM) and domoic acid (DA) analyses and mating experiments (Mating). All strains were characterized through at least one of the following molecular markers: 28S, ITS, 18S and *rbc*L, as indicated in the table.**

| Strain name | Site | Station Coordinates Latitude [°N]-Longitude [°E] | Isolation date | EM | DA | Mating | 28S | ITS | 18S | rbcL |
|---|---|---|---|---|---|---|---|---|---|---|
| 1. B367 | LTER-MC | 40º 48.50’-14º 15.00’ | 31/7/ 2007 | | | | X | | | |
| 2. B366 | LTER-MC | 40º 48.50’-14º 15.00’ | 31/7/ 2007 | | | | X | | | |
| 3. B365 | LTER-MC | 40º 48.50’-14º 15.00’ | 31/7/ 2007 | | | | X | | | |
| 4. B364 | LTER-MC | 40º 48.50’-14º 15.00’ | 31/7/ 2007 | | | | X | | | |
| 5. B359 | LTER-MC | 40º 48.50’-14º 15.00’ | 28/08/2007 | | | | X | | | |
| 6. B363 | LTER-MC | 40º 48.50’-14º 15.00’ | 28/08/2007 | | | | X | | | |
| 7. B361 | LTER-MC | 40º 48.50’-14º 15.00’ | 28/08/2007 | | | | X | | | |
| 8. B358 | LTER-MC | 40º 48.50’-14º 15.00‘ | 28/08/2007 | | | | X | | | |
| 9. B362 | LTER-MC | 40º 48.50’-14º 15.00’ | 28/08/2007 | | | | X | | | |
| 10. MC784_4II_2 | LTER-MC | 40º 48.50’-14º 15.00‘ | 02/10/2007 | | | | X | | | |
| 11. MC784_4II | LTER-MC | 40º 48.50’-14º 15.00‘ | 02/10/2007 | | | | X | | | |
| 12. MC784 B6 | LTER-MC | 40º 48.50’-14º 15.00‘ | 02/10/2007 | | | | X | | | |
| 13. MC784 B2 | LTER-MC | 40º 48.50’-14º 15.00‘ | 02/10/2007 | | | | X | | | |
| 14. MC784 A3 | LTER-MC | 40º 48.50’-14º 15.00’ | 02/10/2007 | | | | X | | | |
| 15. B495 | Ionian Sea | 39º 46.00’-19º 07.06’ | 21/09/2008 | X | X | | X | | | X |
| 16. B853 | Ionian Sea | 39°46.00’-19°07.06’ | 21/09/2008 | X | | | X | X | | X |
| 17. 21ott08strain3 | LTER-MC | 40º 48.50’-14º 15.00’ | 21/10/2008 | | | | X | | | |
| 18. B497 | LTER-MC | 40º 48.50’-14º 15.00’ | 28/07/2009 | X | | | X | | | |
| 19. B498 | LTER-MC | 40º 48.50’-14º 15.00’ | 28/07/2009 | X | | | X | | | |
| 20. B499 | LTER-MC | 40º 48.50’-14º 15.00’ | 28/07/2009 | X | | | X | X | | X |
| 21. B500 | LTER-MC | 40º 48.50’-14º 15.00’ | 28/07/2009 | X | | | X | | | |
| 22. B501 | LTER-MC | 40º 48.50’-14º 15.00’ | 28/07/2009 | X | | | X | X | | X |
| 23. B503 | LTER-MC | 40º 48.50’-14º 15.00’ | 28/07/2009 | X | | | X | | | |
| 24. B504 | LTER-MC | 40º 48.50’-14º 15.00’ | 28/07/2009 | X | | | X | | | |
| 25. B507 | LTER-MC | 40º 48.50’-14º 15.00’ | 28/07/2009 | X | | | X | | | |
| 26. B509 | LTER-MC | 40º 48.50’-14º 15.00’ | 28/07/2009 | X | | | X | X | | X |
| 27. B514 | LTER-MC | 40º 48.50’-14º 15.00’ | 04/08/2009 | X | | | X | | | |
| 28. B522 | LTER-MC | 40º 48.50’-14º 15.00’ | 29/09/2009 | | | | X | | | |
| 29. B523 | LTER-MC | 40º 48.50’-14º 15.00’ | 29/09/2009 | X | | | X | | | |
| 30. B524 | LTER-MC | 40º 48.50’-14º 15.00’ | 29/09/2009 | | X | | X | X | | X |
| 31. B536 | LTER-MC | 40º 48.50’-14º 15.00’ | 29/09/2009 | | | | X | | | |
| 32. B535 | LTER-MC | 40º 48.50’-14º 15.00’ | 29/09/2009 | | | | X | | | |
| 33. B534 | LTER-MC | 40º 48.50’-14º 15.00’ | 29/09/2009 | | | | X | | | |
| 34. B521 | LTER-MC | 40º 48.50’-14º 15.00’ | 29/09/2009 | | | | X | | | |
| 35. B525 | LTER-MC | 40º 48.50’-14º 15.00’ | 06/10/2009 | | | | X | X | | X |

| | | | | | | | | | |
|---|---|---|---|---|---|---|---|---|---|
| 36. B526 | LTER-MC | 40º 48.50’-14º 15.00’ | 06/10/2009 | | | | X | | |
| 37. B527 | LTER-MC | 40º 48.50’-14º 15.00’ | 06/10/2009 | | | | X | | |
| 38. B537 | LTER-MC | 40º 48.50’-14º 15.00’ | 06/10/2009 | | | | X | | |
| 39. B540 | LTER-MC | 40º 48.50’-14º 15.00’ | 04/11/2009 | | | | X | | |
| 40. B541 | LTER-MC | 40º 48.50’-14º 15.00’ | 17/11/2009 | | | | X | | |
| 41. Int-3I-A3 | Mergellina harbour, Gulf of Naples | 40º 49.53’-13º 86.00’ | 05/10/2015 | | | | | X | |
| 42. B631 | Gulf of Naples | 40º 48.50’-14º 15.00’ | NA | | | | | | X |
| 43. Int-3I-A4 | Mergellina harbour, Gulf of Naples | 40º 49.53’-13º 86.00’ | 05/10/2015 | | | | | X | |
| 44. MC1177-D5 | LTER-MC | 40º 48.50’-14º 15.00’ | 10/10/2015 | | | | | X | |
| 45. MC1177-D6 | LTER-MC | 40º 48.50’-14º 15.00’ | 10/10/2015 | | | | | X | |
| 46. MC1177-C6 | LTER-MC | 40º 48.50’-14º 15.00’ | 10/10/2015 | | | | | X | |
| 47. MC1028-B5 | LTER-MC | 40º 48.50’-14º 15.00’ | 05/07/2016 | | | X | | X | |
| 48. MC1028-C3 | LTER-MC | 40º 48.50’-14º 15.00’ | 05/07/2016 | | | X | | X | |
| 49. MC1028-C5 | LTER-MC | 40º 48.50’-14º 15.00’ | 05/07/2016 | | | X | | X | |
| 50. MC1209-A1 | LTER-MC | 40º 48.50’-14º 15.00’ | 12/07/2016 | | | X | | X | |
| 51. MC1209-A2 | LTER-MC | 40º 48.50’-14º 15.00’ | 12/07/2016 | | | X | | X | |
| 52. MC1209-A4 | LTER-MC | 40º 48.50’-14º 15.00’ | 12/07/2016 | | | X | | X | |
| 53. MC1209-B1 | LTER-MC | 40º 48.50’-14º 15.00’ | 12/07/2016 | | | X | | X | |
| 54. MC1209-B3 | LTER-MC | 40º 48.50’-14º 15.00’ | 12/07/2016 | | | X | | X | |
| 55. MC1209-B4 | LTER-MC | 40º 48.50’-14º 15.00’ | 12/07/2016 | | | X | | X | |
| 56. MC1209-B6 | LTER-MC | 40º 48.50’-14º 15.00’ | 12/07/2016 | | | X | | X | |
| 57. MC1209-C1 | LTER-MC | 40º 48.50’-14º 15.00’ | 12/07/2016 | | | X | | X | |
| 58. MC1209-C2 | LTER-MC | 40º 48.50’-14º 15.00’ | 12/07/2016 | | | X | | X | |
| 59. MC1209-C3 | LTER-MC | 40º 48.50’-14º 15.00’ | 12/07/2016 | | | X | | X | |
| 60. MC1209-C3/A | LTER-MC | 40º 48.50’-14º 15.00’ | 12/07/2016 | | | X | | X | |
| 61. MC1209-C4 | LTER-MC | 40º 48.50’-14º 15.00’ | 12/07/2016 | | | X | | X | |
| 62. MC1209-C5 | LTER-MC | 40º 48.50’-14º 15.00’ | 12/07/2016 | | | X | | X | |
| 63. MC1209-D3 | LTER-MC | 40º 48.50’-14º 15.00’ | 12/07/2016 | | | X | | X | |
| 64. MC1209-D4 | LTER-MC | 40º 48.50’-14º 15.00’ | 12/07/2016 | | | X | | X | |

NA=not available.

Table S2: GenBank accession numbers of the strains used for phylogenetic analyses for each molecular marker.

| 18 S | | |
|---|---|---|
| **Species** | **Strain designation** | **GB Accession Number** |
| *Fragilariopsis curta* | Strain 3 | EF140623 |
| *Fragilariopsis cylindrus* | NIES-3887 | LC189151 |
| *P. allochrona* | SZN-B631 | KJ608076 |
| *P. americana* | SKLMP_Sh004 | MG799146 |
| *P. arctica* | RCC2004 | JF794046 |
| *P. arenysensis* | SZN-B593 | XXXXXXX |
| *P. australis* | SPC21 | GU373961 |
| *P. australis* | POMXaus | AM235384 |
| *P. batesiana* | PnTb19 | KP708989 |
| *P. brasiliana* | PnSm07 | KP708990 |
| *P. caciantha* | PnSL05 | KP708992 |
| *P. calliantha* | NWFSC185 | JN091716 |
| *P. circumpora* | PnPd28 | KP708994 |
| *P. cuspidata* | PnPd29 | KP708995 |
| *P. decipiens* | PnKk38 | KP708996 |
| *P. delicatissima* | SZN-B653 | KJ608075 |
| *P. dolorosa* | SZN-B592 | XXXXXXX |
| *P. fraudulenta* | SZN-B670 | KJ608077 |
| *P. fukuyoi* | PnTb25 | KP708997 |
| *P. galaxiae* | SZN-B606 | KJ608078 |
| *P. galaxiae* | SZN-B617 | KJ608079 |
| *P. granii* | RCC2008 | JN934671 |
| *P. heimii* | NWFSC205 | JN091727 |
| *P. kodamae* | PnPd31 | KP709000 |
| *P. lineola* | NWFSC188 | JN091717 |
| *P. lundholmiae* | PnTb21 | KP709001 |
| *P. mannii* | SZN-B640 | KJ608080 |

| | | |
|---|---|---|
| *P. micropora* | PnKk14 | KP709003 |
| *P. multiseries* | NparI | AM235380 |
| *P. multiseries* | Tka2 | U18241 |
| *P. multistriata* | VF2.3 | XXXXXXX |
| *P. pseudodelicatissima* | SZN-B656 | KJ608082 |
| *P. pungens* | PnKd05 | KP709004 |
| *P. sabit* | PnPd82 | KP709005 |
| *P. subcurvata* | UNC1409 | KX253952 |
| *P. turgidula* | NWFSC220 | FJ222752 |

Table S2, ctd.

| 28 S | | |
|---|---|---|
| **Species** | **Strain designation** | **GB Accession Number** |
| *Cylindrotheca fusiformis* | UTEX2083 | AF417665 |
| *P. abrensis* | Ner-J2 | KP172231 |
| *P. allochrona* | SZN-B509 | KC801042 |
| *P. allochrona* | SZN-B495 | XXXXXXX |
| *P. allochrona* | SZN-B507 | KC801041 |
| *P. americana* | CV2 | U41390 |
| *P. arctica* | RCC2002 | JQ995416 |
| *P. arenysensis* | AL-24 | DQ813811 |
| *P. arenysensis* | SZN-B33 | AF416758 |
| *P. australis* | OM1 | AF417651 |
| *P. batesiana* | PnTb19 | KC147534 |
| *P. bipertita* | Pnmi04 | KR021334 |
| *P. brasiliana* | Brasil 8 | AF469672 |
| *P. calliantha* | AL112 | DQ813841 |
| *P. chiniana* | MC3011 | MN128956 |
| *P. circumpora* | PnSb58 | KC147533 |
| *P. cuspidata* | AL-17 | DQ813809 |
| *P. decipiens* | Mex13 | EF506608 |
| *P. delicatissima* | AL22 | DQ813810 |
| *P. delicatissima* | CV3 | U41391 |
| *P. dolorosa* | AL59 | DQ813813 |
| *P. fraudulenta* | Limens1 | AF417647 |
| *P. fryxelliana* | NWFSC241 | JN050296 |
| *P. fukuyoi* | PnTb25 | KC147535 |
| *P. galaxiae* | Mex23 | AY081136 |
| *P. granii* | RCC2008 | JQ995421 |
| *P. hallegraeffii* | CTD44 2 | MF044022 |
| *P. hallegraeffii* | CTD44 3 | MF044024 |
| *P. hasleana* | NWFSC252 | JN050298 |
| *P. inflatula* | No7 | AF417639 |
| *P. kodamae* | PnPd36 | KF482045 |

| | | |
|---|---|---|
| *P. kodamae* | PnPd26 | KF482042 |
| *P. kodamae* | PnPd39 | KF482046 |
| *P. limii* | Pnmi16 | KR021343 |
| *P. linea* | ICMB-156 | FJ489633 |
| *P. lineola* | NWFSC188 | JN050300 |
| *P. lundholmiae* | PnTb10 | KC147538 |
| *P. mannii* | AL101 | DQ813814 |
| *P. micropora* | VPB.B3 | AF417649 |
| *P. multiseries* | NWFSC 011 | AF440772 |
| *P. multistriata* | SZN-B27 | AF416753 |
| *P. nanaoensis* | MC4188 | MG787875 |
| *P. plurisecta* | Ner-A1 | KP172228 |
| *P. pseudodelicatissima* | AL-15 | DQ813808 |
| *P. pungens* | CV4 | U41262 |
| *P. pungens* | NWFSC32 | AF440776 |
| *P. qiana* | MC3007 | MN128952 |
| *P. sabit* | Ps283 | KP288514 |
| *P. seriata* | Nissum3 | AF417652 |
| *P. subcurvata* | CCMP1431 | HQ396851 |
| *P. subfraudulenta* | rensubfrau | AF417646 |
| *P. subpacifica* | RdA8 | AF417642 |
| *P. turgidula* | NWFSC255 | JN050301 |
| *P. turgiduloides* | 124C | EF531709 |
| *P. simulans* | MC281 | MF374774 |

Table S2, ctd.

| ITS | | |
|---|---|---|
| **Species** | **Strain designation** | **GB Accession Number** |
| *Fragilariopsis kerguelensis* | 4-20 | EF660061 |
| *P. abrensis* | Ner-J2 | KC409108 |
| *P. allochrona* | SZN-B853 | XXXXXXX |
| *P. allochrona* | SZN-B509 | XXXXXXX |
| *P. allochrona* | SZN-B524 | XXXXXXX |
| *P. allochrona* | SZN-B525 | XXXXXXX |
| *P. americana* | Kervel | EU523099 |
| *P. arctica* | P2F2N2 | KT589421 |
| *P. arenysensis* | MexA | DQ329211 |
| *P. arenysensis* | AL-24 | DQ813830 |
| *P. arenysensis* | Ner-D1 | GQ228393 |
| *P. australis* | OEM1 | AY257842 |
| *P. batesiana* | PnTb19 | KC147514 |
| *P. bipertita* | Pnmi04 | KR021318 |
| *P. brasiliana* | Pnkk31 | JN252429 |
| *P. bucculenta* | L1.3 | MH376341 |
| *P. caciantha* | AL-56 | DQ813834 |
| *P. calliantha* | AL-112 | DQ813841 |
| *P. cf. subpacifica* | RdA8 | AY257860 |
| *P. chiniana* | MC3011 | MK411963 |
| *P. circumpora* | PnSb58 | JN252430 |
| *P. cuspidata* | AL-17 | DQ813827 |
| *P. decipiens* | GranCan4-1 | DQ336157 |
| *P. decipiens* | Mex13 | DQ336156 |
| *P. delicatissima* | Tasm 10 | AY257848 |
| *P. delicatissima* | AL-22 | DQ813829 |
| *P. delicatissima* | Laesoe5 | AY257849 |
| *P. dolorosa* | AL-59 | DQ813835 |
| *P. fraudulenta* | NWFSC 200 | FJ222755 |
| *P. fraudulenta* | Limens1 | AY257840 |
| *P. fryxelliana* | NWFSC 241 | JN050288 |

| | | |
|---|---|---|
| *P. fukuyoi* | PnTb25 | KC147516 |
| *P. galaxiae* | Mex23 | AY257850 |
| *P. galaxiae* | Sydney4 | DQ336158 |
| *P. granii* | UBC100 | EU051654 |
| *P. hainanensis* | MC3099 | MW042679 |
| *P. hainanensis* | MC3098 | MW042678 |
| *P. hallegraeffii* | CTD44 2 | MF044023 |
| *P. hallegraeffii* | CTD44 3 | MF044025 |
| *P. hasleana* | NWFSC 186 | JN050282 |
| *P. inflatula* | no7 | DQ329204 |
| *P. kodamae* | Pnmi66 | KR021307 |
| *P. kodamae* | Pnmi67 | KR021308 |
| *P. limii* | Pnmi16 | KR021311 |
| *P. lineola* | NWFSC 188 | JN091756 |
| *P. lundholmiae* | PnTb10 | KC147523 |
| *P. mannii* | AL-101 | DQ813839 |
| *P. micropora* | no16 | DQ329209 |
| *P. micropora* | VPB-B3 | AY257847 |
| *P. multiseries* | mu3 | AY257844 |
| *P. multistriata* | KoreaA | AY257843 |
| *P. nanaoensis* | MC4188 | MG787881 |
| *P. obtusa* | T5 | DQ062667 |
| *P. plurisecta* | Ner-F1 | KC409089 |
| *P. pseudodelicatissima* | AL-15 | DQ813826 |
| *P. pungens* | PnMt45 | HQ111412 |
| *P. pungens var. aveirensis* | P-24 | AY257845 |
| *P. qiana* | MC3007 | MK412843 |
| *P. qiana* | MC989 | MK412841 |
| *P. qiana* | MC991 | MK412842 |
| *P. sabit* | PnPd57 | KM400610 |
| *P. seriata* | Nissum3 | AY257841 |
| *P. simulans* | MC281 | MF374769 |
| *P. subcurvata* | 1-F | DQ329205 |
| *P. taiwanensis* | MC5109 | MW042680 |

| | | |
|---|---|---|
| *P. turgidula* | NWFSC220 | JN091764 |
| *P. turgiduloides* | 3-19 | AY257839 |

Table S2, ctd.

| *rbc*L | | |
|---|---|---|
| **Species** | **Strain designation** | **GB Accession Number** |
| *Cylindrotheca sp.* | N1 | M59080 |
| *P. allochrona* | SZN-B499 | as SZN-B507 |
| *P. allochrona* | SZN-B501 | as SZN-B507 |
| *P. allochrona* | SZN-B524 | as SZN-B507 |
| *P. allochrona* | SZN-B525 | as SZN-B507 |
| *P. allochrona* | SZN-485 | as SZN-B507 |
| *P. allochrona* | SZN-583 | as SZN-B507 |
| *P. allochrona* | SZN-B509 | as SZN-B507 |
| *P. allochrona* | SZN-B507 | KC801037 |
| *P. americana* | FBJun06.6 | EF423504 |
| *P. arctica* | RCC2002 | KT808257 |
| *P. arenysensis* | AL64 | DQ813823 |
| *P. arenysensis* | AY11 | EF423516 |
| *P. arenysensis* | SZN-B487 | KC801036 |
| *P. arenysensis* | AL24 | DQ813819 |
| *P. caciantha* | AL56 | DQ813821 |
| *P. calliantha* | Al117 | DQ813825 |
| *P. cuspidata* | AL28 | DQ813820 |
| *P. delicatissima* | AL22 | DQ813818 |
| *P. dolorosa* | AL59 | DQ813822 |
| *P. fraudulenta* | AL75 | EF520333 |
| *P. fraudulenta* | AL50 | EF423502 |
| *P. fraudulenta* | BB19 | EF423503 |
| *P. fryxelliana* | NWFSC 241 | JN050302 |
| *P. galaxiae* | SM3 | EF423512 |
| *P. galaxiae* | AL8 | EF423515 |
| *P. galaxiae* | SM54 | EF423514 |
| *P. hasleana* | NWFSC 186 | JN050304 |
| *P. mannii* | AL101 | DQ813824 |
| *P. multiseries* | NWFSC 316 | KC801040 |
| *P. multistriata* | 279 | EF520337 |

| | | |
|---|---|---|
| *P. multistriata* | 19A | EF423505 |
| *P. multistriata* | DD4 | EF520336 |
| *P. pseudodelicatissima* | AL15 | DQ813817 |
| *P. pseudodelicatissima* | SZN-B317 | KC801039 |
| *P. pungens* | Na213 | FM207548 |
| *P. pungens* | FBA2A11 | EF423507 |
| *P. turgiduloides* | 7A1 | EF423508 |

Table S3: Morphometric data of *P. allochrona* and morphologically related species. Average and standard deviation in brackets, when available.

| | Reference | Valve shape | Width (µm) | Length (µm) | Central nodule | Striae in 10 µm | Fibulae in 10 µm | Poroids in 1 µm | Rows of poroids | Band striae in 10 µm |
|---|---|---|---|---|---|---|---|---|---|---|
| ***P. allochrona*** | This study | Lanceolate | 1.4-2.1<br>(1.8 ± 0.2) | 32-84* | + | 34-44<br>(40.3 ± 2.1) | 20-26<br>(23.3 ± 1.5) | 10-12<br>(10.7 ± 1.6) | 2 | 46-50<br>(47.7 ± 1.5) |
| **(as *P.* cf. *arenysensis*)** | Giulietti et al. 2021 | Linear | 1.5-2.3<br>(1.9 ± 0.2 | 29.1-50.6<br>(38.5 ± 5.2) | + | 36-42<br>(38.3 ± 1.6) | 16-36<br>(21.5 ± 2.2) | 8-12<br>(10.1 ± 0.9) | 2 | 42-52<br>(46.0 ± 2.3) |
| ***P. arenysensis*** | Quijano-Scheggia et al. 2009 | Lanceolate | 1.6–2.5 | 22–84** | + | 34–43 | 20–26 | 7–12 | 2 | 40–50 |
| | Ajani et al. 2013 | Lanceolate, symmetrical | 1.8-2.7<br>(2.1 ± 0.2) | 33.6-45.8<br>(39.8 ± 5.3) | + | 38-45<br>(40.4 ± 1.5) | 20-26<br>(22.8 ± 2.2) | 8-11<br>(9.4 ± 0.9) | 2 | 40-46<br>(42 ± 2.8) |
| ***P. bucculenta*** | Gai et al. 2018 | Lanceolate | 2.7-3.6<br>(3.0 ± 0.3) | 19-31<br>(24.9 ± 3.6) | + | 28-35<br>(31.4 ± 1.7) | 16-21<br>(18.4 ± 1.2) | 5-7.5<br>(6.7 ± 0.6) | 1-2 | 38-39<br>(38 ± 0.6) |
| ***P. chiniana*** | Huang et al 2019 | Lanceolate | 2.3-2.6<br>(2.4 ± 0.1) | 42-58<br>(49.7 ± 1.5) | + | 30-34<br>(32.8 ± 1.3) | 17-22<br>(19.3 ± 1.8) | 4-6<br>(5 ± 1) | 1-2 | 38-40 |
| ***P. decipiens*** | Lundholm et al. 2006 | Lanceolate | 1.4-2.4<br>(1.9 ± 0.3) | 29-64 | + | 41-46<br>(43.2 ± 1.2) | 20-26<br>(24.0 ± 1.4) | 9-13<br>(11.4 ± 1.2) | 2 | 48-55<br>(51.8 ± 1.7) |
| | Teng et al. 2015 | Lanceolate, symmetrical | 1.7-2.0 | 41.8-49.1 | + | 43-47 | 22-26 | 8-13 | 2 | 48-54 |
| ***P. delicatissima*** | Lundholm et al. 2006 | Lanceolate | 1.5-2.0<br>(1.8 ± 0.2) | 19-94* | + | 35-40<br>(36.8 ± 1.5) | 19-26<br>(21.4 ± 1.6) | 8-12<br>(10.1 ± 1.2) | 2 | 43-48<br>(44. ± 1.6) |
| ***P. dolorosa*** | Lundholm et al. 2006 | Lanceolate, asymmetrical | 2.5-3.0<br>(2.6 ± 0.2) | 30-59 | + | 30-36<br>(34.5 ± 1.4) | 18-22<br>(20.0 ± 1.0) | 5-8<br>(6.6 ± 0.8) | 1-2 | 40-44<br>(42.0 ± 1.4) |
| | Lim et al 2012 | nd | 1.8-2.1<br>(2.0 ± 0.2) | 42.4-43.0<br>(42.7 ± 0.3) | + | 35-37<br>(36.2 ± 0.8) | 21-22<br>(21.6 ± 0.5) | 5-6<br>(5.8 ± 0.5) | 1 | nd |
| ***P. hainanensis*** | Chen et al. 2021 | Lanceolate | 1.8-2.0 | 23-45 | + | 30-34 | 17-22 | 4-8 | 2 | 41-42 |
| ***P. hallegraeffii*** | Ajani et al. 2018 | Lanceolate, asymmetrical | 1.9-3.1 | 25.6-55.4 | + | 34-40 | 16-22 | 6-8 | 1-2 | 43-56 |
| ***P. micropora*** | Priisholm et al. 2002 | Lanceolate | 1.3-2.0 | 31-57 | - | 41-46 | 21-29 | 9-12 | 2 | 48-54 |
| | Ajani et al. 2013 | Lanceolate, symmetrical | 1.8-2.3<br>(2.0 ± 0.1) | 33.1-36.0<br>(34.9 ± 0.9) | - | 42-50<br>(45.9 ± 2.9) | 23-30<br>(27.1 ± 2.7) | 9-13 | 2 | 54-60<br>(59.3 ± 2.1) |
| ***P. prolongatoides*** | Almandoz et al. 2008 | Lanceolate | 1.5-2.6 | 20-85 | + | 29-33 | 16-21 | 10-13 | 2-3 | nd |
| ***P. sabit*** | Teng et al. 2015 | Falcate, asymmetrical | 1.4-2.6 | 22.5-69.0* | + | 38-45 | 17-26 | 7-13 | 2 | 50-58 |
| ***P. turgiduloides*** | Almandoz et al. 2008 | Lanceolate | 2-2.9 | 81-126 | + | 18-24 | 10-14 | 7-10 | 1-2 | nd |
| ***P. turgidula*** | Almandoz et al. 2008 | Lanceolate | 2.3–2.5 | 41–79 | + | 24–28 | 15–18 | 7–9 | 2-3 | nd |
| ***P. yuensis*** | Dong et al. 2020 | Lanceolate | 1.8-2.5 | 37-44 | + | 38-43 | 19-24 | 6-7 | 1-2 | 43-48 |

*: maximum length measured on the initial cell; **: maximum length from the initial cell length from Amato et al. (2005, as *P. delicatissima*).

***Table S4:*** **Results of cross experiments between couples of different strains of *P. allochrona* and between 4 of them with 2 *P. arenysensis* strains of different mating types.** Strain names abbreviated retaining only the final part of their name as reported in Table S1. By convention, female sex was attributed to the strains to which zygotes were attached, identified by their size in EM images from crosses between strains with cells of different sizes (Fig. 2 D), and then extrapolated to all strains sexually incompatible with those ones.

| | *P. allochrona* | | | | | | | | | | | | | | | | | | *P. arenysensis* | |
|---|---|---|---|---|---|---|---|---|---|---|---|---|---|---|---|---|---|---|---|---|
| | **8-B5** ♀ | **8-C3** ♂ | **8-C5** ♀ | **9-A1** ♀ | **9-A2** ♂ | **9-A4** ♀ | **9-B1** ♀ | **9-B3** ♂ | **9-B4** ♂ | **9-B6** ♀ | **9-C1** ♂ | **9-C2** ♂ | **9-C3** ♂ | **9-C3A** ♀ | **9-C4** ♀ | **9-C5** ♀ | **9-D3** ♂ | **9-D4** ♀ | **BB16** ♂ | **CM63** ♀ |
| **8-B5♀** | | + (2) | - (2) | - | +(2) | - | - | + | + | - | + | + | + | - | - | - | + | - | - | - |
| **8-C3♂** | | | + (2) | + | - (2) | + | + | - | - | + | - | - | - | + | + | + | - | + | - | - |
| **8-C5♀** | | | | - | +(2) | - | - | + | + | - | + | + | + | - | - | - | + | - | - | - |
| **9-A1♀** | | | | | + | - | - | + | + | - | + | + | + | - | - | - | + | - | | |
| **9-A2♂** | | | | | | + | + | - | - | + | - | - | - | + | + | + | - | + | - | - |
| **9-A4♀** | | | | | | | - | + | + | - | + | + | + | - | - | - | + | - | | |
| **9-B1♀** | | | | | | | | + | + | - | + | + | + | - | - | - | + | - | | |
| **9-B3♂** | | | | | | | | | - | + | - | - | - | + | + | + | - | + | | |
| **9-B4♂** | | | | | | | | | | + | - | - | - | + (*) | + (*) | + (*) | - | + | | |
| **9-B6♀** | | | | | | | | | | | + | + | + | - | - | - | + | - | | |
| **9-C1♂** | | | | | | | | | | | | - | - | + | + | + | - | + | | |
| **9-C2♂** | | | | | | | | | | | | | - | + | + | + | - | + | | |
| **9-C3♂** | | | | | | | | | | | | | | + | + | + | - | + | | |
| **9-C3A♀** | | | | | | | | | | | | | | | - | - | + (*) | - | | |
| **9-C4♀** | | | | | | | | | | | | | | | | - | + | - | | |
| **9-C5♀** | | | | | | | | | | | | | | | | | + | - | | |
| **9-D3♂** | | | | | | | | | | | | | | | | | | + | | |
| **9-D4♀** | | | | | | | | | | | | | | | | | | | | |
| **BB16♂** | | | | | | | | | | | | | | | | | | | | + |
| **CM63♀** | | | | | | | | | | | | | | | | | | | | |

+: sexual reproduction observed
–: sexual reproduction not observed
(2): replicated crosses
*: crosses used for electron microscopy observations
♀: female mating types
♂: male mating types
dark grey boxes: crosses not performed

Table S5: Estimates of net evolutionary divergence between species of the *P. delicatissima*-complex closest to *P. allochrona*, using a Maximum Composite Likelihood model. Standard error estimates (italics, above the diagonal) were estimated through 100 bootstrap replicates.

| | *P. allochrona* | *P. arenysensis* | *P. delicatissima* | *P. dolorosa* | *P. decipiens* | *P. micropora* |
|---|---|---|---|---|---|---|
| *P. allochrona* | | *0.010* | *0.011* | *0.028* | *0.023* | *0.013* |
| *P. arenysensis* | 0.042 | | *0.012* | *0.030* | *0.023* | *0.009* |
| *P. delicatissima* | 0.090 | 0.090 | | *0.026* | *0.019* | *0.007* |
| *P. dolorosa* | 0.251 | 0.283 | 0.246 | | *0.025* | *0.025* |
| *P. decipiens* | 0.189 | 0.205 | 0.163 | 0.258 | | *0.016* |
| *P. micropora* | 0.082 | 0.045 | 0.024 | 0.234 | 0.133 | |
